# Low-order MLKL oligomers induce irregular membrane defects in necroptosis

**DOI:** 10.64898/2026.10.21.760398

**Authors:** Veronica Martinez-Osorio, Qingrong Zhang, Silvia Zanna, Jonas Aufdermauer, Uris Ros, Ana J. Garcia-Saez

## Abstract

MLKL is a key executioner of necroptosis, a lytic form of regulated cell death associated with inflammatory pathologies. Upon activation by RIPK3-mediated phosphorylation, MLKL oligomerizes and translocates to the plasma membrane to induce permeabilization through yet unknown mechanisms. By imaging single particles of MLKL at the plasma membrane of necroptotic cells, here we show that, upon activation, MLKL transitions from a diffuse cytosolic distribution to discrete, low-mobility assemblies before cell bursting. This is accompanied by oligomerization into a mixture of species ranging from monomers up to hexamers. We directly visualize MLKL-induced membrane damage using atomic force microscopy imaging in supported bilayers, which reveals the formation of irregular membrane openings whose edges are only partially decorated with protein. Our findings support a mechanism in which low-order self-assemblies of MLKL disrupt plasma membrane integrity and stabilize membrane defect edges without organizing into defined pore structures, in contrast to membrane permeabilization in other forms of cell death. This reframes necroptosis execution and dissects membrane insertion of MLKL oligomers for potential targeting.

## INTRODUCTION

Necroptosis is an inflammatory form of regulated cell death that has been associated to a wide range of human pathologies, including cancer (Seifert *et al*., 2016; Cho, 2018; Ando *et al*., 2020; Martens *et al*., 2021; Yu *et al*., 2022), neurodegeneration (Oñate *et al*., 2020; Moujalled, Strasser and Liddell, 2021; Richard and Mousa, 2022), and cardiovascular disease (Puylaert *et al*., 2022). In its canonical form, necroptosis is initiated by activation of the Tumor Necrosis Factor (TNF) engagement under conditions of caspase inhibition, which triggers assembly of a supramolecular complex known as the necrosome. The necrosome is initially assembled through the interaction of the Receptor-Interacting Serine/Threonine Kinase 1 and 3 (RIPK1 and RIPK3). Subsequently, the pseudokinase Mixed Lineage Kinase domain-Like protein (MLKL) is recruited to the complex, where it becomes activated, oligomerizes, and translocates to the plasma membrane, where it promotes permeabilization, ultimately leading to cell death (Li *et al*., 2021).

Despite intense research, the mechanism by which MLKL mediates plasma membrane damage and cell death execution remains poorly understood. In particular, the properties of the MLKL assemblies responsible for membrane disruption and the nature of the membrane alterations induced by these MLKL oligomers are still unclear. Two broad mechanistic models have been proposed: one in which MLKL organizes into defined, protein-lined pore structures, and one in which it permeabilizes membranes through partial insertion and accumulation in the lipid bilayer pointing out a detergent or carpet-like mechanism (Flores-Romero, Ros and Garcia-Saez, 2020). Proposed structures include nanopores (Ros *et al*., 2017), cation channels (Xia *et al*., 2016; Huang *et al*., 2017), hotspot-like accumulations (Samson *et al*., 2020), fiber-like polymers (Liu *et al*., 2024), and amyloid-like assemblies (Liu *et al*., 2017), yet no consensus exists on whether a defined protein pore forms and, if so, what its architecture is.

The precise stoichiometry and structure of the MLKL oligomers responsible for necroptosis execution remain a matter of debate. Evidence suggests that tetramers are more prevalent in human cells (Tanzer *et al*., 2015, 2016), whereas trimers predominate in mice (Chen *et al*., 2014); however, higher-order oligomers have also been reported (Huang *et al*., 2017; Petrie, Hildebrand and Murphy, 2017; Garcia *et al*., 2021). Our current knowledge of the oligomeric structure and stoichiometry of MLKL is derived primarily from Western blotting (WB), negative-stain electron microscopy, and small-angle X-ray scattering (SAXS). However, these methods characterize the dominant species in solution and cannot resolve heterogeneous oligomeric distributions or directly visualize MLKL architecture at the plasma membrane. Structural modelling with AlphaFold and ColabFold Multimer was used to propose that human MLKL (hMLKL) adopts a dimerization-driven tetrameric active conformation in solution, via brace extension and release of the four-helix bundle (4HB) domain (Meng *et al*., 2023). Nevertheless, whether these species predominate at the membrane of living necroptotic cells, and their dynamic properties at the membrane are still lacking at single-molecule resolution.

Here, we address this gap by combining single-molecule Total Internal Reflection Fluorescence (TIRF) microscopy with atomic force microscopy (AFM) to visualize MLKL at the plasma membrane of living necroptotic cells and membrane mimetic systems at single-molecule resolution. TIRF allowed us to obtain a detailed *in situ* view of MLKL oligomerization and dynamics at the plasma membrane. By comparing the properties of necroptosis-proficient MLKL isoforms with those of its inactive splicing MLKL isoforms (Ros *et al*., 2025), we identify unique key features associated with the MLKL mechanism of action during necroptosis execution. We show that active, but not inactive MLKL isoforms transition from diffuse cytosolic distribution to low-mobility plasma membrane-bound particles during necroptosis. Brightness and photobleaching analyses reveal that these particles are heterogeneous in size, suggesting that MLKL can assemble at the plasma membrane into complexes of varying stoichiometry. Importantly, *in vitro* liposome permeabilization and AFM on supported lipid bilayers (SLBs) experiments reveal that MLKL disrupts lipid membranes by forming stable and irregular openings only partially decorated by the protein, which we directly visualize. These observations are consistent with both proteins and lipids participating in the formation of membrane defects, in which MLKL stabilizes membrane edges. Together, our findings uncover key molecular steps in the mechanism of necroptosis execution by MLKL that may guide drug design targeting MLKL at the plasma membrane for clinical applications.

## RESULTS

### Comparison of active/inactive MLKL isoforms reveals determinants of necroptotic function in humans and mice

In previous work, we reported an inactive isoform of MLKL (mMLKL1 in mouse, hMLKL0 in human) that differs from the necroptosis-proficient MLKL isoform (mMLKL2 in mouse, hMLKL1 in human) in a few amino acid insertions at the C-terminal helix (Ros *et al*., 2025). We reasoned that comparing the dynamics and assembly of both isoforms would enable dissecting the functional features of active, necroptosis-proficient MLKL. To enable single-molecule visualization of MLKL at the plasma membrane, we employed a Tet-On system to express the active (hMLKL1, mMLKL2) and necroptosis-deficient (hMLKL0, mMLKL1) isoforms as C-terminal EGFP fusions in *MLKL^−/−^* HT-29 and *Mlkl^−/−^* MDF cells (Figure 1A,B). A non-dimerizing GFP variant (K260A) was used to avoid potential oligomerization artifacts (Zacharias, Violin and Newton, 2002). To select comparable expression levels across conditions while maintaining sufficiently low expression for single-particle detection, we first quantified EGFP fluorescence intensity over time using the Incucyte system (Supplementary Figure 1A,B). Based on this, we defined MLKL-GFP expression conditions (doxycycline (Dox), a Tet-On system activator, at 1 μg/mL for 1 h) and characterized the membrane permeabilization kinetics following TSZ (necroptosis induction; TNF, Smac-mimetic LCL161, and caspase inhibitor z-VAD-fmk) treatment, monitoring DRAQ7 (membrane permeabilization marker) uptake with the Incucyte (Figure 1C,D). As expected, cells expressing active MLKL isoforms displayed efficient DRAQ7 internalization, whereas cells expressing the inactive variants did not.

**Figure 1.**
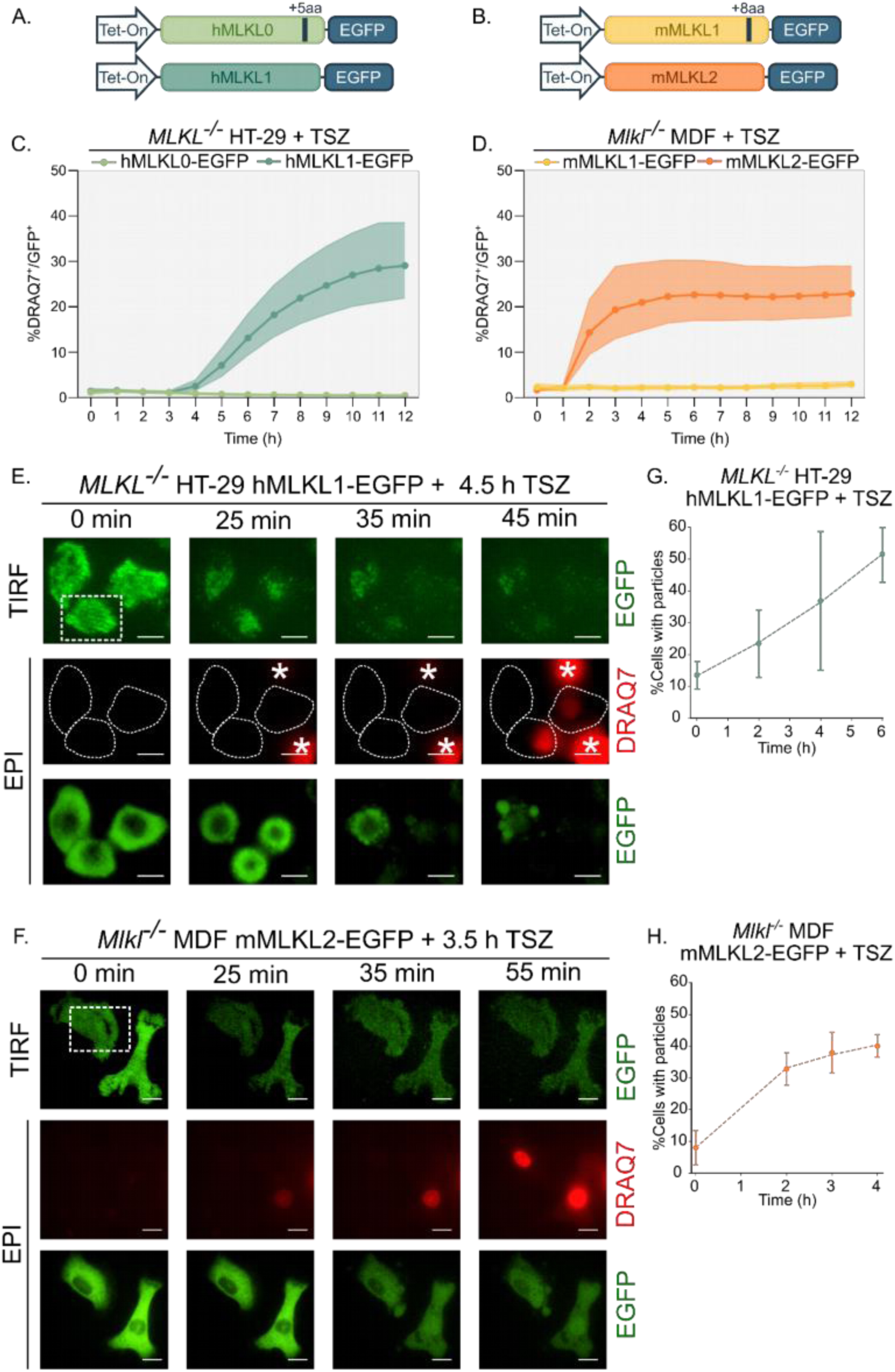
Active MLKL reorganizes into plasma membrane particles before necroptotic membrane rupture. (A, B) Schematic of the Tet-On expression constructs for human (A) and mouse (B) MLKL isoforms C-terminally tagged with EGFP. hMLKL0 carries a +5 aa insert; mMLKL1 carries a +8 aa insert relative to the respective active isoform. (C, D) Kinetics of plasma membrane permeabilization measured by DRAQ7 uptake in *MLKL^−/−^* HT-29 human cells (C) and *Mlkl^−/−^* MDF mouse cells (D) expressing the constructs in (A) and (B), treated with doxycycline (1 µg/mL, 1 h) followed by TSZ (TNF 30 ng/mL, LCL-161 10 µM, Z-VAD-FMK 10 µM). Data are mean ± SD; n = 3 independent experiments, each performed in triplicate wells. (E, F) Representative time-lapse TIRF and epifluorescence (EPI) images of *MLKL^−/−^* HT-29 cells expressing hMLKL1-EGFP (E) and *Mlkl^−/−^* MDF cells expressing mMLKL2-EGFP (F) during necroptosis induction (TSZ treatment). Imaging was performed 4.5–5.5 h (human) or 3.5–4.5 h (mouse) after TSZ addition. TIRF row: plasma membrane-restricted EGFP signal. EPI rows: DRAQ7 (membrane integrity) and EGFP (total cellular). The cell shapes at imaging time 0 are indicated in dashed lines on the EPI-DRAQ7 Images. Asterisks indicate other dead cells that entered the field while imaging. Dashed white boxes indicate regions containing MLKL-EGFP particles, shown at higher magnification in Supplementary figure 1G (E) and 1H (F). Scale bar, 10 µm. (G, H) Percentage of cells displaying plasma membrane MLKL-EGFP particles over time after TSZ treatment, in *MLKL^−/−^* HT-29 hMLKL1-EGFP cells (G) and *Mlkl^−/−^* MDF mMLKL2-EGFP cells (H). Data are mean ± SD; n = 2-3 independent experiments. The temporal increase in particle-bearing cells closely parallels DRAQ7 uptake in (C) and (D), indicating that MLKL particle formation precedes but anticipates membrane permeabilization. TSZ, TNF + LCL-161 + Z-VAD-FMK necroptosis-inducing cocktail; UT, untreated; TIRF, total internal reflection fluorescence microscopy; EPI, epifluorescence microscopy; DRAQ7, far-red DNA intercalator used as membrane permeabilization marker; DOX, doxycycline.

mMLKL1 has impaired phosphorylation by RIPK3 at S345 site and is able to form basal complexes/oligomers that appear in the cytosol and membrane fractions (Ros *et al*., 2025). However, whether a similar mechanism applied to hMLKL0 remained unclear. To address this, we performed WB from cell lysates expressing the human isoforms. Similar to mMLKL1, inactive hMLKL0 was unable to undergo phosphorylation at S358 upon TSZ treatment (Supplementary figure 1C). In contrast to mMLKL1, hMLKL0 did not show enrichment in the membrane fraction or formation of oligomeric complexes, neither under basal nor necroptotic conditions, providing a more direct mechanistic basis for its functional inactivity (Supplementary figure 1D). Altogether, characterization of the Tet-On system validated the use of active and inactive MLKL isoforms as a robust framework for defining features associated with MLKL necroptotic function and for gaining mechanistic insights into differences in isoform activity.

### Active MLKL shifts from a diffuse to a discrete particle distribution at the plasma membrane during necroptosis

We then applied single-molecule TIRF to investigate the dynamic behavior and stoichiometry of MLKL-EGFP versions directly at the membrane in real time. TIRF allows *in situ* visualization of single particles at or near the plasma membrane by providing adequate sectioning by restricting fluorophore excitation to 100–200 nm above the coverslip (Fish, 2022). Unlike bulk biochemical methods, which report ensemble-averaged properties, single-molecule TIRF can resolve individual particle stoichiometry and lateral mobility directly at the membrane in real time. In these experiments, we acquired images at defined time windows after TSZ treatment. We selected the 4.5–5.5 h in human cells and the 3.5–4.5 h in mouse cells (Supplementary figure 1E,F) as time windows that allowed robust detection of necroptotic cell death at similar expression levels between isoforms (Figure 1C,D). Simultaneous acquisition of DRAQ7 fluorescence in epifluorescence mode allowed us to monitor plasma membrane permeabilization while tracking MLKL-EGFP particles by TIRF at a single-cell level. In cells expressing the active hMLKL1/mMLKL2 isoforms, individual MLKL-EGFP particles appeared at the plasma membrane before DRAQ7 entry (Figure 1E,F and Supplementary figure 1G,H). The fraction of cells displaying MLKL particles increased over time (Figure 1G,H), closely paralleling the rise in DRAQ7-positive cells (Figure 1C,D). The temporal relation between particle formation and imminent membrane rupture suggests that MLKL particle assembly at the plasma membrane is a mechanistic step in the process of necroptosis execution.

MLKL-EGFP could be detected as single particles at or in close proximity to the plasma membrane across all conditions, especially in the case of mouse MLKL. We observed that the active isoforms hMLKL1-EGFP and mMLKL2-EGFP formed prominent, brighter particles under TSZ treatment, accompanied by the disappearance of the diffuse MLKL-EGFP signal (Figure 2A,B). This transition was not detected in cells expressing the inactive isoforms. To better characterize the distribution of MLKL within cells, we performed a phenotypic classification of MLKL-EGFP signal patterns across all conditions. Based on our observations, we defined three categories: (I) diffuse signal, (II) diffuse signal with single particles, and (III) single particles without diffuse signal (Figure 2C and Supplementary figure 2A,B). In the case of human MLKL, cells expressing the active MLKL isoforms (hMLKL1) primarily displayed a diffuse signal (type I) under untreated conditions that shifted to distinct MLKL-EGFP brighter particles without a diffuse signal (type III) upon TSZ treatment (Figure 2D,E). A similar transition was observed for mMLKL2, although it presented a more mixed distribution in untreated cells, which included a fraction of cells with diffuse signal and single particles (type II). In contrast, cells expressing the inactive isoforms showed diffuse signal coexisting with single particles (type II) both in basal and necroptotic conditions. These observations indicate that necroptosis induction drives the redistribution from diffuse to membrane-associated particles for active MLKL isoforms, but not for necroptosis-deficient ones. This feature thus appears to be specifically associated with MLKL activation, likely playing a causal role in necroptotic cell death.

**Figure 2.**
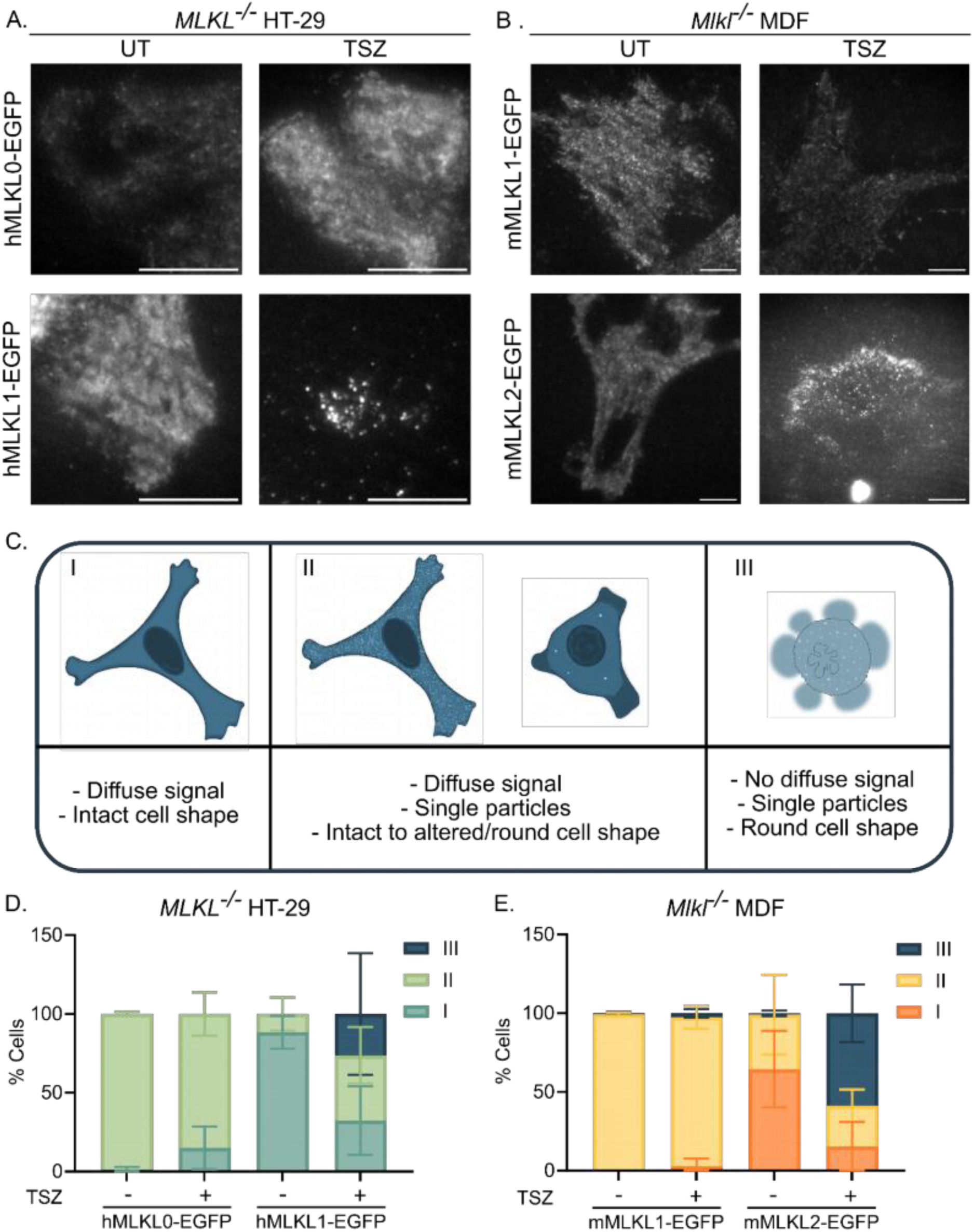
MLKL activation triggers isoform-specific redistribution from diffuse cytosolic to discrete plasma membrane particles. (A, B) Representative TIRF images of *MLKL^−/−^* HT-29 cells (A) and *Mlkl^−/−^* MDF cells (B) expressing the indicated EGFP-tagged MLKL isoforms under untreated (UT) or TSZ-treated conditions. In both cell types, active isoforms (hMLKL1, mMLKL2) form bright discrete particles under TSZ, whereas inactive isoforms (hMLKL0, mMLKL1) retain a diffuse signal. Scale bar, 10 µm. (C) Schematic classification of the three EGFP signal distribution phenotypes used for quantification: Type I, uniform diffuse signal with intact cell morphology; Type II, diffuse signal coexisting with single particles and variable cell rounding; Type III, discrete single particles without diffuse signal, typically in rounded/blebbing, permeabilized cells. (D, E) Frequency of each phenotype in *MLKL^−/−^* HT-29 cells (D) and *Mlkl^−/−^* MDF cells (E), under basal and TSZ-treated conditions. Only cells expressing the active MLKL isoforms show a marked increase in Type III cells upon TSZ treatment. Data are mean ± SD from n = 3 independent experiments; ∼60 cells per condition per experiment. Note that SD can extend beyond 100% in stacked bar graphs due to between-experiment variability in individual category proportions. TSZ, necroptosis-inducing cocktail (TNF + LCL-161 + Z-VAD-FMK); UT, untreated; TIRF, total internal reflection fluorescence microscopy.

### Active MLKL forms confined, long-lived membrane assemblies consistent with stable membrane engagement

To characterize the mobility of MLKL-EGFP particles at or near the plasma membrane, we performed single-particle tracking on short TIRF time-lapse movies (5.5 s, 25 ms/frame). The individual MLKL-EGFP particles were detected and tracked using the TrackIt software (Kuhn *et al*., 2021) and employing the developer-defined u-track algorithm based on local intensity maxima. Analysis of particle dynamics under necroptotic conditions revealed the active isoform of human MLKL (hMLKL1) displayed a significant increase in the time-averaged track length of detected particles, indicative of an increased residence time at the plasma membrane (Figure 3A,B and Supplementary figure 3A). This increase was accompanied by a reduction in the mean jump distance between consecutive frames (Figure 3C-H and Supplementary figure 3B), which is indicative of reduced mobility. A similar behavior, but less pronounced, was observed for active mouse MLKL (mMLKL2).

**Figure 3.**
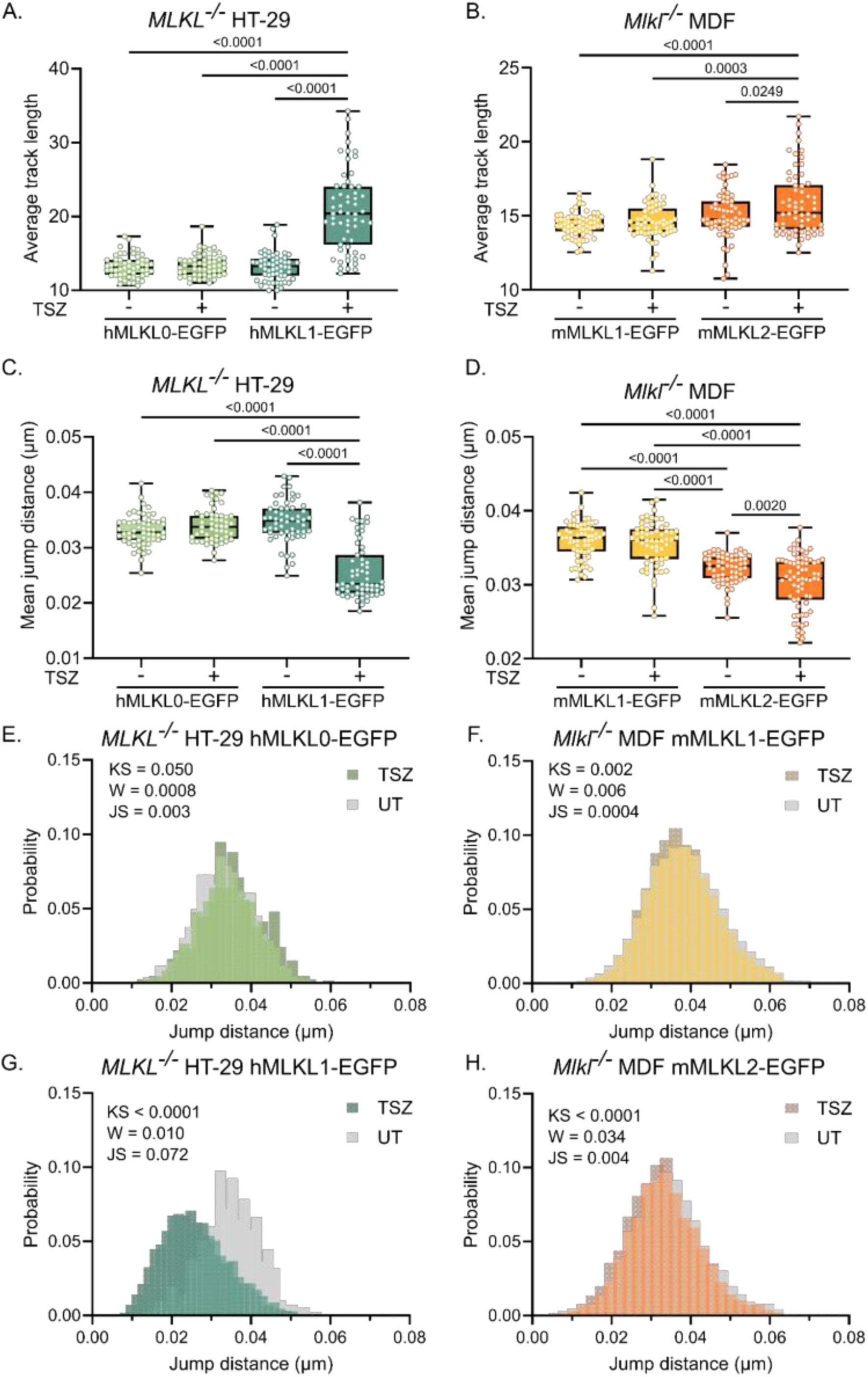
Active MLKL particles increase their residence time and reduce their mobility at the plasma membrane during necroptosis. (A, B) Per-movie average single particle track length (frames) of MLKL-EGFP particles detected by single-particle tracking TIRF microscopy, in *MLKL^−/−^* HT-29 cells (A) and *Mlkl^−/−^* MDF cells (B) expressing the indicated isoforms, under basal and TSZ-treated conditions. Each data point represents one movie (field of view). Box: median and interquartile range; whiskers: 5th–95th percentile. P-values from Tukey’s multiple comparisons test following one-way ANOVA; n = ∼ 60 movies from 3 independent experiments. 1 frame = 25 ms. (C, D) Per-movie mean jump distance (µm) between consecutive frames for the same cells as in (A) and (B). A reduction in mean jump distance reflects decreased lateral mobility. Statistics as in (A, B). (E–H) Probability distributions of individual particle jump distances under UT and TSZ conditions for each isoform: hMLKL0-EGFP in HT-29 (E), mMLKL1-EGFP in MDF (F), hMLKL1-EGFP in HT-29 (G), and mMLKL2-EGFP in MDF (H). Inset p-values indicate pairwise comparisons between UT and TSZ distributions using three distribution-distance metrics: KS, two-sample Kolmogorov–Smirnov test statistic; W, Wasserstein distance; JS, Jensen–Shannon divergence. Note that for the inactive isoforms (E, F) the distributions are statistically indistinguishable, whereas active isoforms (G, H) show a marked shift towards lower jump distance under TSZ treatment, reflecting reduced mobility. TSZ, necroptosis-inducing cocktail; UT, untreated; TIRF, total internal reflection fluorescence microscopy.

In line with this, mean squared displacement (MSD) analysis showed a clear decrease in the diffusion coefficient, from 0.012±0.007 μm^2^/s to 0.006±0.006 μm^2^/s, upon activation of hMLKL1 under TSZ treatment (Supplementary figure 3C-F). A smaller, yet significant, shift in the distribution of diffusion coefficients was detected for mMLKL2, with an average decrease from 0.010±0.006 μm^2^/s to 0.009±0.006 μm^2^/s. In contrast, this value remained invariable for the inactive isoforms (hMLKL0/mMLKL1) under untreated and treated conditions. This range of diffusion coefficient is typical of integral membrane proteins, for example Kv2.1 (∼0.03 µm²/s, clustered) (Tamkun, O’Connell and Rolig, 2007), EGFR (0.04–0.08 µm²/s) (Lagerholm *et al*., 2017), and TrkB/p75 (∼0.01 µm²/s) (Chein, Perlson and Roichman, 2019), and is substantially below the lower bound of GPI-anchored proteins (≥0.05 µm²/s) (Saha *et al*., 2015), which do not traverse the bilayer. Taken together, these data are consistent with stable membrane engagement of active MLKL under necroptotic conditions, potentially involving partial or full insertion rather than a superficial interaction with the membrane.

### Active MLKL exists as a heterogeneous population ranging from monomers to hexamers at the plasma membrane during necroptosis

To determine the molecular stoichiometry of MLKL-EGFP particles at the plasma membrane, we performed a single-particle fluorescent brightness analysis. In this approach, the number of molecules per particle is inferred from the fluorescence intensity measured in photon counts relative to a reference of known stoichiometry. The intensity of a monomeric EGFP was determined from a lysate of cells expressing soluble EGFP under the exact conditions used for imaging mouse and human cells (Supplementary figure 4A,B) and used to calculate the expected fluorescence intensity of MLKL-EGFP oligomers (Supplementary figure 4C,D). These theoretical values were then used to fit the fluorescence intensity distribution obtained for the single particles detected from the images using a multi-Gaussian model, in which each Gaussian component corresponds to a specific oligomeric species. These values were further corrected considering that only 80% of EGFP molecules are fluorescent due to misfolding events and the proportion of MLKL molecules allocated in each oligomeric species is estimated (Figure 4A) (Schmidt, 1996; Subburaj *et al*., 2015). After performing the brightness analysis for human and mouse cells expressing the different MLKL isoforms (Figure 4B-I) we found that hMLKL0 and mMLKL1 molecules appeared predominantly as monomers, with a minor fraction of dimers, and that this distribution remained unchanged upon TSZ treatment (Figure 4B,C,F,G, and Supplementary figure 4E,F). In contrast, hMLKL1 and, to a lesser extent, mMLKL2 showed a marked increase in oligomeric species (Supplementary figure 4G-J), switching from monomers/dimers under basal conditions to a broader distribution also including tetramers. A small proportion of hexamers was also detected specifically for hMLKL1 (Figure 4D,E and Supplementary figure 4G).

**Figure 4.**
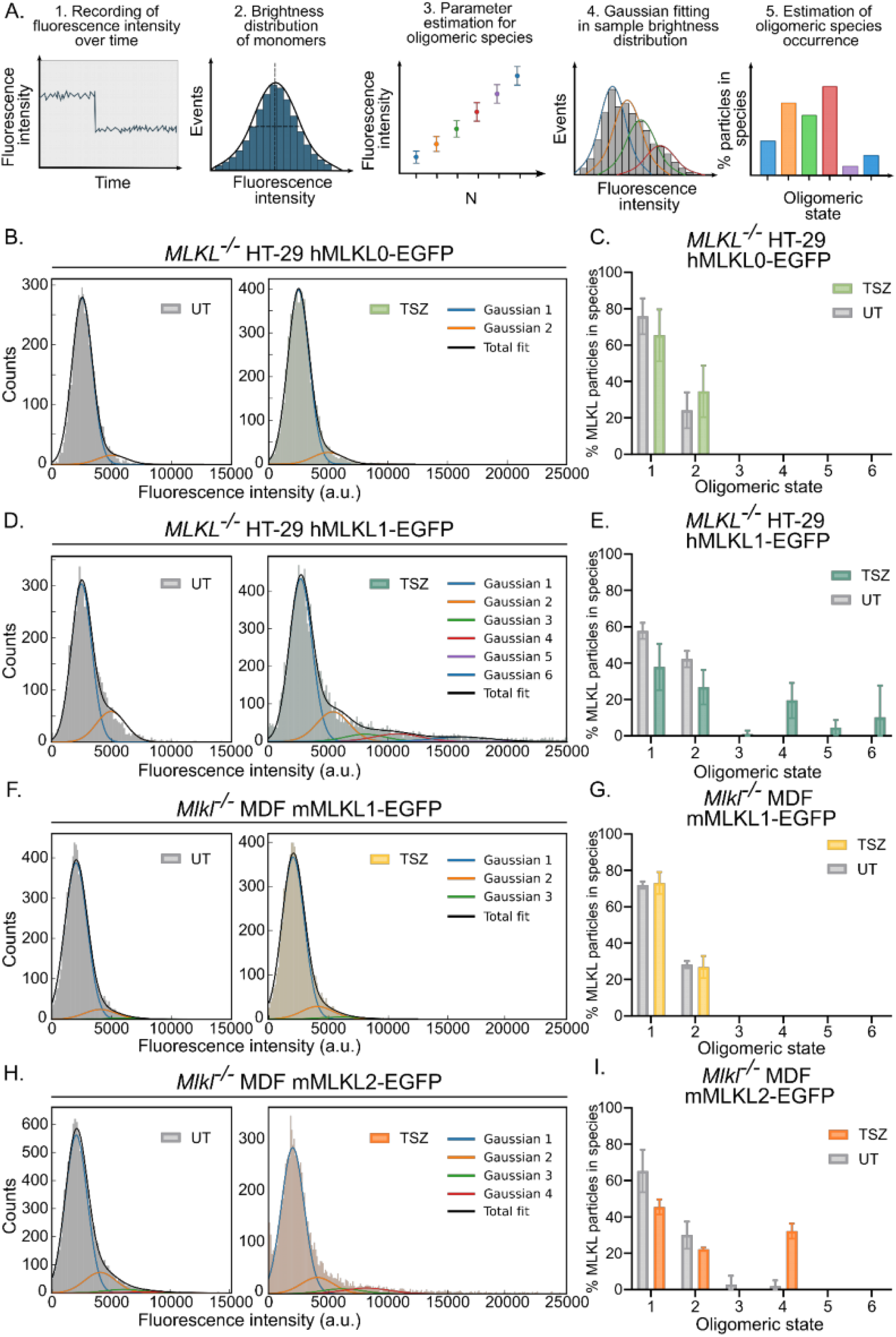
Active MLKL assembles into a mixture of oligomers, from monomers to hexamers, at the plasma membrane during necroptosis. (A) Schematic of the single-particle fluorescence brightness analysis workflow: (1) Fluorescence intensity of each particle is recorded over time under continuous illumination. (2) Photobleaching steps in the calibration sample (soluble mEGFP) identify monomers and are used to callibrate the monomer intensity distribution. (3) From the monomer parameters, the expected intensities for oligomeric species (dimer, trimer, etc.) are calculated. (4) The brightness distribution of MLKL-EGFP particles in the sample is fitted by multi-Gaussian decomposition, where each Gaussian component corresponds to one oligomeric species. (5) The fractional occurrence of each species is calculated and corrected for EGFP labelling efficiency (80%, accounting for fluorescent-protein misfolding). The percentage of MLKL molecules allocated in each species is calculated by weighting the corrected occurrence considering the respective N-mer fraction. (B–I) Brightness distributions (left panels, UT; right panels, TSZ) and corresponding fractional occurrence of oligomeric states (bar graphs) for: hMLKL0-EGFP in HT-29 (B, C); hMLKL1-EGFP in HT-29 (D, E); mMLKL1-EGFP in MDF (F, G); mMLKL2-EGFP in MDF (H, I). In each brightness histogram, individual Gaussian components (colored lines) correspond to monomers through the maximum detected oligomeric state; the black line shows the total fit. Note that the near-empty appearance of UT histogram panels for mouse isoforms (F, H left) reflects the low basal particle density in untreated MDF cells (see also Supplementary figure 3F). Active isoforms (hMLKL1, mMLKL2) display a marked redistribution toward higher-order species upon TSZ treatment. Data are mean ± SD; data from 3 independent experiments (n=3). TSZ, necroptosis-inducing cocktail; UT, untreated; EGFP, enhanced green fluorescent protein.

To validate these findings by an alternative approach, we employed subunit photobleaching analysis (Subburaj *et al*., 2015). In this method, the time evolution of single particle photobleaching is collected until all particles are photobleached, and the number of fluorophores per particle is estimated by counting discrete photobleaching steps (Figure 5A, and Supplementary figure 5A,B). Since photobleaching is a stochastic process, each step corresponds to the photobleaching of one fluorophore. We performed subunit photobleaching analysis on the hMLKL1-GFP and mMLKL2-GFP single particles formed in TSZ-treated cells (Figure 5B), which revealed that they exist at the plasma membrane as a distribution of states spanning from monomers to hexamers (Figure 5C). Analysis of photobleaching step amplitudes revealed comparable intensity distributions across steps, supporting a linear relationship between fluorescence intensity and molecule number and discarding potential homoFRET effects (Supplementary figure 5C,D). Together, these results indicate that active MLKL does not adopt a single defined stoichiometry at the plasma membrane, but instead appears as a heterogeneous population of monomers and low-order oligomeric assemblies, supporting a mechanism of assembly upon activation at the plasma membrane.

**Figure 5.**
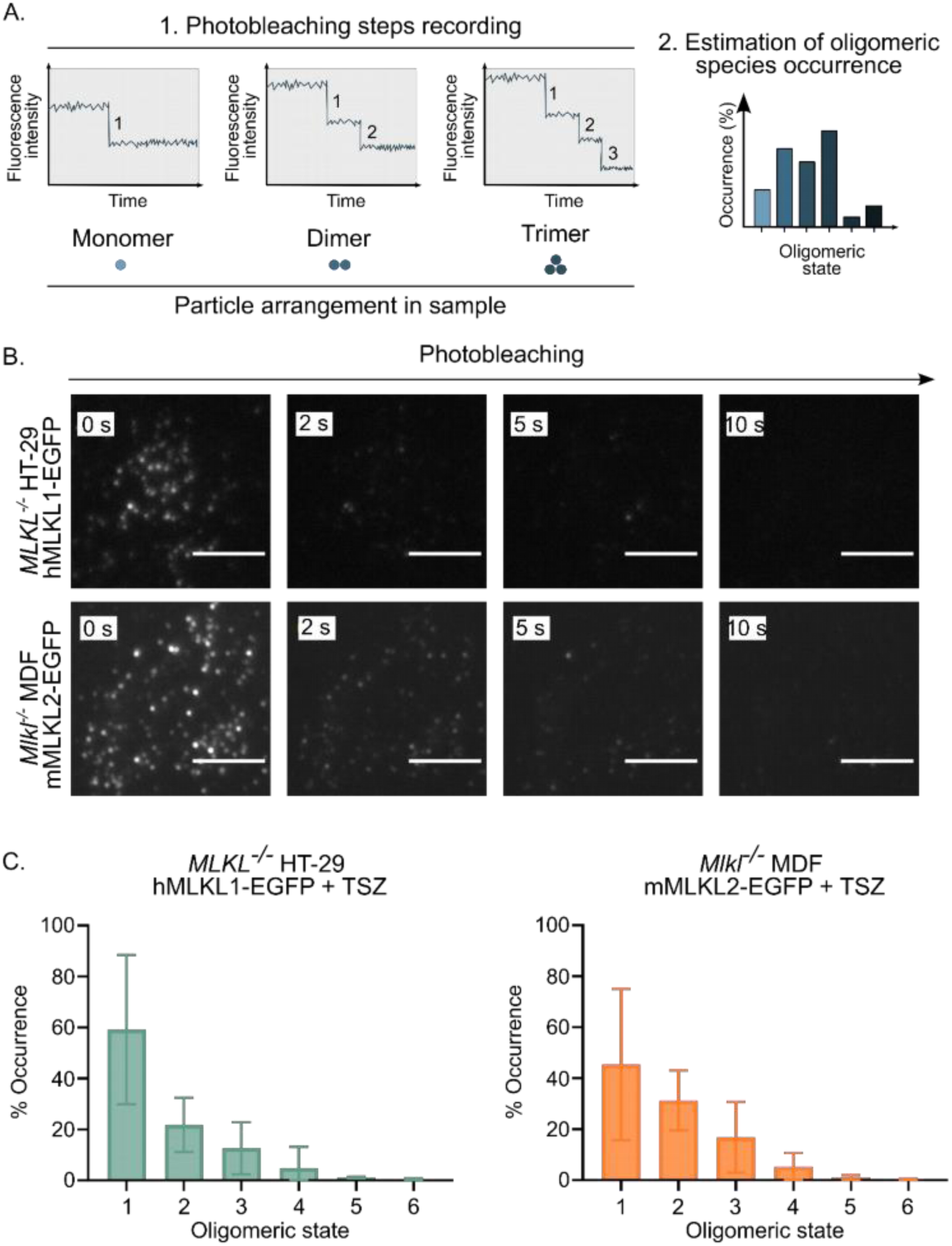
Subunit photobleaching analysis confirms that active MLKL forms a heterogeneous oligomeric population dominated by monomers and low-order assemblies. (A) Principle of subunit photobleaching analysis stoichiometry. Particles are imaged under continuous illumination until all fluorophores are bleached. The number of discrete intensity steps to baseline corresponds to the number of EGFP molecules per particle, allowing direct counting of oligomeric state. (B) Representative TIRF image sequences showing progressive photobleaching of hMLKL1-EGFP (top) and mMLKL2-EGFP (bottom) particles in cells treated with TSZ. Images shown at 0, 2, 5, and 10 s. Scale bar, 10 µm. (C) Fractional occurrence of each oligomeric state (monomers through hexamers) determined by step counting for hMLKL1-EGFP in *MLKL^−/−^* HT-29 cells (left) and mMLKL2-EGFP in *Mlkl^−/−^* MDF cells (right) under TSZ treatment. Monomers represent the most abundant species (∼60% in human, ∼45% in mouse), with diminishing fractions of dimers, trimers, and higher-order assemblies. The distribution is consistent with the brightness analysis in Figure 4 (D, E, H, I), validating the two orthogonal stoichiometric approaches. Data are mean ± SD; data from 3 independent experiments. Inactive isoforms (hMLKL0, mMLKL1) were excluded from this analysis owing to insufficient particle density for reliable step counting. TSZ, necroptosis-inducing cocktail; TIRF, total internal reflection fluorescence microscopy; EGFP, enhanced green fluorescent protein.

### MLKL induces stable, irregular, partially protein-lined membrane openings in lipid bilayers

To gain mechanistic insight into how MLKL’s interaction with membranes causes damage, we implemented an *in vitro* reconstituted system made of recombinant MLKL and model membranes. We used the monomeric fraction of the purified hMLKL1 recombinant protein expressed in insect cells and vesicles mimicking different inner plasma membrane compositions (Supplementary figure 6A,B). The carboxyfluorescein leakage assay using large unilamellar vesicles (LUVs) enabled quantification of MLKL membrane-permeabilizing activity in bulk across a range of conditions, whereas uptake of externally added fluorescent dyes into giant unilamellar vesicles (GUVs) enabled visualization and tracking of MLKL activity at the single-vesicle level.

In line with previous studies (Wang *et al*., 2014; Petrie *et al*., 2018; Ros *et al*., 2025), we found that MLKL mediated membrane permeabilization in the absence of any additional protein, in a broad range of membrane compositions at nanomolar concentrations. Incubation of increasing concentrations of recombinant hMLKL1 with LUVs led to a robust dose-response release of the dye (Figure 6A). hMLKL1 also altered LUVs’ morphology, reflected in a broader radius distribution of the particles (Supplementary figure 6C), suggesting potential vesicle clustering. Incubation of hMLKL1 with GUVs also resulted in efficient membrane permeabilization (Figure 6B and Supplementary figure 6D), reflected by rapid dye influx kinetics (Figure 6C). Dye influx kinetics into single vesicles were fitted using a one-phase association model. The permeabilization rate constant was K = 0.039 ± 0.017 s⁻¹, corresponding to a time constant of 36.4 s (half-time = 25.2 s). The initial dye influx rate, calculated as K×Span, was 3.3 normalized fluorescence units s⁻¹. Remarkably, MLKL activity did not compromise vesicle integrity, discarding a detergent-like mechanism (Sandre, O., Moreaux, L. and Brochard-Wyart, 1999; Sudbrack *et al*., 2011; Kulma and Anderluh, 2021). Most vesicles were fully permeabilized, supporting an all-or-none mechanism of permeabilization instead of a graded one. We also tested the stability of the permeabilized state by the sequential addition of two different dyes of similar size to the sample (Figure 6D,E). The efficient entry of the second dye into the GUVs indicated that permeabilization was stable, at least for the time interval between the addition of the dyes (75 min).

**Figure 6.**
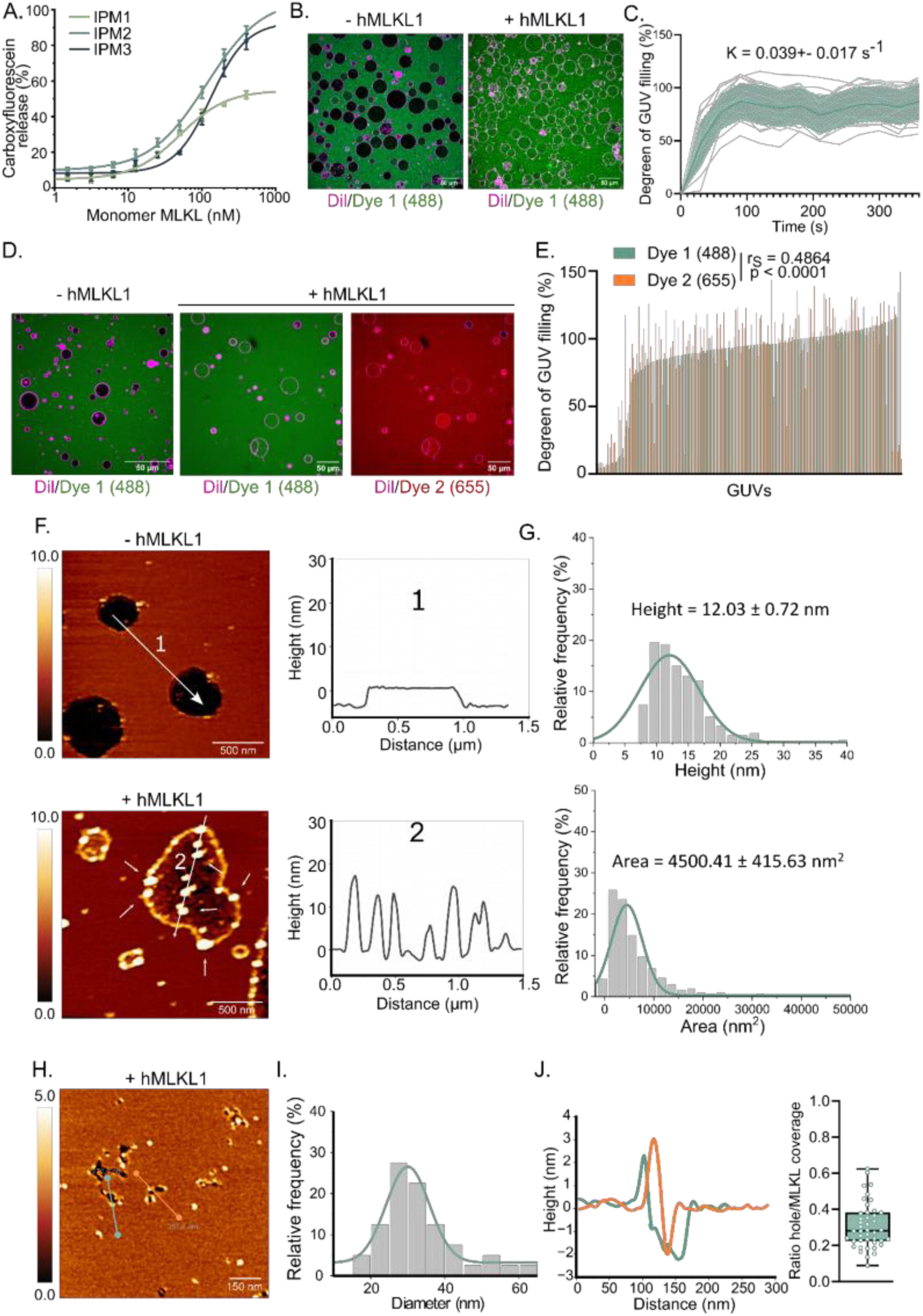
Recombinant hMLKL1 permeabilizes membranes by forming irregular, partially protein-lined openings. (A) Dose–response liposome permeabilization assay. Carboxyfluorescein-loaded large unilamellar vesicles (LUVs) mimicking three inner plasma membrane lipid compositions (IPM1: PI; IPM2: PI(4,5)P₂; IPM3: PI(3,4,5)P₃; full compositions in Supplementary figure 6A) were incubated with increasing concentrations of monomeric recombinant hMLKL1. Dye release is expressed as percentage of maximum (detergent lysis). Data are mean ± SD; n = 3 independent preparations. (B) Confocal fluorescence images of giant unilamellar vesicles (GUVs) before and after incubation with hMLKL1. GUV membrane is labelled with DiI (magenta); internalized dye (Dye 1, 488 nm, green) indicates permeabilization. Scale bar, 50 µm. (C) Mean fluorescence intensity of Dye 1 (488 nm) inside individual GUVs from (B), measured over time after hMLKL1 addition. Rapid dye influx kinetics (K=0.039+-0.017 s^-1^) without vesicle disruption indicate membrane permeabilization rather than detergent-like solubilization. Gray lines represent individual GUVs and colored line represents the mean ± SD (n = 13). (D) Two-dye sequential GUVs permeabilization assay. GUVs were incubated with hMLKL1; Dye 1 (488 nm, green) and Dye 2 (655 nm, red) were added sequentially 75 min apart. Scale bar, 50 µm. (E) Paired quantification of GUVs filling by Dye 1 and Dye 2 (n = 179 GUVs). The positive Spearman correlation (rs = 0.49, p < 0.0001) indicates that pores formed by hMLKL1 remain stable for at least 75 min. (F) AFM height images of supported lipid bilayers (SLBs) in the absence (top) and presence (bottom) of hMLKL1. Line profiles (numbered 1 and 2) show smooth bilayer surface without protein (1) and protein cluster accumulation at bilayer edges (2, indicated by white arrows). (G) Height and area distribution of hMLKL1 clusters detected by AFM from (F). (H) AFM images of SLBs pre-incubated with hMLKL1 prior to bilayer formation. White arrows indicate irregular membrane openings lined by hMLKL1. (I) Diameter frequency distribution of the observed membrane defects; mean diameter ≈ 30 nm, but no uniform pore size is detected (n = 43 pores from 8 SLBs). Scale bar: 150 nm. (J) Quantification of the fraction of each pore perimeter decorated by hMLKL1 (hole/MLKL coverage ratio). Left: height profiles of indicated lines in (H). Right: box and whisker plot (median ± IQR; whiskers, 5th–95th percentile). SLB, supported lipid bilayer; GUV, giant unilamellar vesicle; LUV, large unilamellar vesicle; AFM, atomic force microscopy; IPM, inner plasma membrane; DiI, 1,1′-dioctadecyl-3,3,3′,3′-tetramethylindocarbocyanine perchlorate.

To directly visualize the effect of hMLKL1 on lipid membranes at nanoscale resolution, we performed AFM on SLBs. Here, we implemented two complementary experimental workflows. In the first, SLBs were formed in the absence of hMLKL1, and once assembled on the mica, the protein was added to the buffer. While control SLBs presented smooth surfaces with some defects largely devoid of protein density signal in the surrounding area, samples incubated with hMLKL1 showed an accumulation of the protein on the surface, but especially at the edges of the defects, forming cluster-like structures (Figure 6F,G and Supplementary figure 6E). These results indicate that hMLKL1 exhibits a preference for binding to the edges of membrane bilayers.

In a second approach, liposomes were first incubated with hMLKL1 to form proteoliposomes, which were then deposited onto the mica. hMLKL1 induced the formation of irregular openings on the membrane surface (Figure 6H) and localized preferentially at the edges of these defects, consistent with our previous observations (Figure 6F). Quantitative analysis showed that these membrane defects presented an average diameter of approximately 30 nm (Figure 6I) but lacked a uniform shape or defined architecture, in line with low line tension at the edge. Critically, quantification of pore perimeter decoration revealed that every defect examined was only partially lined by hMLKL1 (coverage ratio < 1.0 in all cases; Figure 6J), a structural hallmark that distinguishes these openings from protein-lined pores, in which the protein completely encircles the opening. Taken together, these *in vitro* data demonstrate that hMLKL1 permeabilizes membranes by inducing stable and irregular openings that are partially lined by protein molecules, instead of defined protein pores.

## DISCUSSION

MLKL is the key executioner of plasma membrane permeabilization and lytic cell death in necroptosis, a function exerted through mechanisms that have so far remained elusive. In this study, we establish three interconnected features of MLKL action: a phosphorylation-triggered membrane binding step that converts diffuse MLKL into low-mobility assemblies, accompanied by heterogeneous oligomeric organization at the membrane surface and the opening of irregular, partially protein-lined membrane defects. Together, these support a model in which the plasma membrane functions both as the assembly platform and as an active structural participant in the permeabilization process.

The shift of MLKL from a diffuse intracellular localization to low-mobility oligomeric assemblies with longer residency at the plasma membrane is likely governed by RIPK3-dependent phosphorylation, as only the isoforms capable of undergoing this post-translational modification displayed this transition. This dynamic reorganization is consistent with membrane insertion and oligomerization based on the decreased diffusion coefficient measured for active MLKL particles. Therefore, longer membrane engagement of MLKL oligomers emerges as a critical step in necroptosis, likely enabling the local accumulation of active MLKL molecules required for reaching a critical threshold of membrane permeabilization. Inactive human and mouse MLKL isoforms are not RIPK3-phosphorylated, do not stably anchor at the plasma membrane, and remain as monomers and dimers even upon necroptosis induction, underscoring the differential behavior of naturally existing MLKL isoforms (Ros *et al*., 2025).

Previous studies have proposed that MLKL oligomerizes at the necrosome, then de-engages and translocates as an oligomer to the plasma membrane (Samson *et al*., 2020; Garnish *et al*., 2021). However, this model does not account for our observations of monomeric MLKL at the membrane, also evident in bulk WB analysis in other studies (Cai *et al*., 2014; Huang *et al*., 2017). Our findings support an alternative assembly mechanism so that two non-mutually exclusive scenarios can be envisioned: MLKL oligomers assemble at the necrosome in the cytosol and subsequently translocate to the plasma membrane, or alternatively, membrane-associated MLKL molecules progressively assemble at the membrane after translocation. While our data do not formally exclude either possibility, the heterogeneous oligomeric MLKL distribution at the plasma membrane is more consistent with the latter scenario.

The heterogeneous oligomers formed by active MLKL at the plasma membrane consist predominantly of low-order assemblies. To date, most evidence has supported the existence of trimers (Cai *et al*., 2014), tetramers (Petrie *et al*., 2018; Meng *et al*., 2023), octamers (Huang *et al*., 2017), and even higher-order assemblies, including hotspots (Samson *et al*., 2020) and polymer-like structures (Liu *et al*., 2017, 2024). Our results in living cells suggest that monomers, together with lower-order oligomers, such as trimers and tetramers, may represent the most stable MLKL species during necroptotic conditions. Importantly, our data are consistent in the human and mouse systems, supporting a conserved mechanism underlying MLKL activation, membrane engagement, and oligomerization.

Unlike gasdermins, which form defined oligomeric arcs and rings (Mulvihill *et al*., 2018), and BAX/BAK, which assemble into larger heterogeneous clusters containing even hundreds of molecules (Subburaj *et al*., 2015; Große *et al*., 2016; Salvador-gallego *et al*., 2016), or NINJ1, which organizes into high-order fibrillar structures (Degen *et al*., 2023; Wang *et al*., 2025), we find that low-order assemblies of MLKL can promote structurally irregular openings lined by both protein and lipid. This places MLKL in a distinct mechanistic category among pore-forming executioners of regulated cell death. The partial edge decoration observed by AFM is a structural signature similar to the toroidal lipid-pore model proposed for amphipathic peptides (Yang *et al*., 2001; Zemel, Fattal and Ben-shaul, 2003) and for BAX and BAK (Qian *et al*., 2008; Bleicken *et al*., 2018). By accumulating at the edges of these openings, protein molecules stabilize the open state of the pore. Our observations of partially decorated membrane defects, together with the stable membrane permeabilization maintaining overall vesicle integrity, are also consistent with protein-lipid openings rather than a detergent-like effect (Sandre, O., Moreaux, L. and Brochard-Wyart, 1999; Sudbrack *et al*., 2011; Kulma and Anderluh, 2021). Yet, in contrast to BAX and BAK pores observed by AFM (Salvador-gallego *et al*., 2016), MLKL-induced membrane defects are irregular in shape, suggesting a local disruption of the lipid bilayer that becomes stabilized by MLKL molecules, rather than a membrane-tension driven opening and line-tension driven organization of a lipidic pore (Flores-Romero, Ros and Garcia-Saez, 2020).

Recombinant hMLKL1 is intrinsically membrane-active, inducing permeabilization in model membranes on its own, as a minimal machinery, consistent with previous reports (Wang *et al*., 2014; Petrie *et al*., 2018; Ros *et al*., 2025). However, we cannot exclude the possibility of MLKL interacting with other molecular actors in the cellular context. For example, SIGLEC12 is an adhesion protein proposed to mediate plasma membrane permeabilization downstream of MLKL during necroptosis in human cells (Noh *et al*., 2026). Also important is the proposed interplay between MLKL and the ESCRT (Gong *et al*., 2017; Yoon *et al*., 2017; Fan *et al*., 2019) membrane repair machinery that can limit the extent of membrane permeabilization and likely the plasma membrane residency of MLKL particles. The fact that MLKL forms stable yet irregular membrane defects has broader physiological implications, as the extent of plasma membrane permeabilization and lysis would depend on reaching a critical amount of membrane damage rather than on their stability.

Altogether, our data support a model in which, upon necroptosis induction, active MLKL assembles into a mixture of low-order oligomers inserted into the plasma membrane, which facilitates assembly. MLKL oligomers promote the opening of irregular membrane defects whose edges are only partially lined by protein. These structurally heterogeneous, low-coverage membrane openings represent a mechanistically distinctive mode of regulated plasma membrane permeabilization: one that sets necroptosis apart from pyroptosis, apoptosis, and NINJ1-induced plasma membrane rupture.

### Limitations of the study

Our cellular system conserved the main features of MLKL-induced necroptosis, but we cannot discard the possibility that the use of fluorescently tagged, exogenously expressed MLKL may not fully recapitulate the behavior of endogenous MLKL. Although our imaging approaches enabled the characterization of MLKL dynamics, stoichiometry, and membrane association with high spatial and temporal resolution, they do not allow the direct identification of the oligomeric species responsible for membrane permeabilization. Likewise, the methodologies employed here lack the structural resolution required to determine the ultrastructure of membrane-associated MLKL assemblies. As a result, the relationship between oligomer architecture, stoichiometry, and membrane-disrupting activity remains to be fully established. In addition, while important for establishing minimal function and as a chemically controlled environment to study MLKL activity, our *in vitro* membrane systems do not incorporate the full complexity of the cellular environment, and relevant factors may be missing. Future studies combining structural and functional approaches in model membranes and in cells will be required to directly link individual MLKL oligomeric species with their membrane-permeabilizing activity and to understand their regulation.

## MATERIALS AND METHODS

### Cell culture

Tet-on MDFs were cultured in low-glucose Dulbecco’s Modified Eagle Medium (DMEM, Sigma-Aldrich, Germany) supplemented with 10% Fetal Bovine Serum (FBS) and 1% penicillin-streptomycin (ThermoFisher, Germany), and grown in a humidified incubator containing 5% CO_2_ at 37°C. The cells were passaged every 3 days.

For all imaging experiments, cells were seeded at a density of 1.5 x 10^4^ cells/well in an 8-well glass-bottom chamber (cat: 80827, IBIDI), previously coated with 0.01 % poly-L-lysine. The following day, cells were treated with 1 μg/mL DOX for 1 h prior to induction of cell death by necroptosis.

### Induction of necroptosis

Human and mouse cells were subjected to a treatment with a cocktail containing TNF (30 ng/mL), LCL-161 (10 µM), and Z-VAD-FMK (10 μM) in normal culture media. Incubation times with this treatment vary depending on the downstream experiments described in the following sections.

### IncuCyte experiments

The IncuCyte bioimaging platform (Essen, UK) was used to track EGFP levels and membrane permeabilization by DRAQ7 staining in cells subjected to necroptosis treatment. Experiments were carried out using a 96-well plate format. The percentage of DRAQ7-positive cells was calculated on the GFP-positive population. Due to a significant difference in the cell-boundary detection between the types of cells, we used the basic analysis module of the IncuCyte software for human cells HT29 and the cell-by-cell analysis module for the mouse MDF cells.

### Western blot

The day after seeding, cells were treated with 1 μg/mL DOX for 4 h, followed by the necroptotic cocktail treatment described above for 2 to 4 h (a variation of the incubation times is described specifically for oligomerization and membrane translocation experiments).

After treatment, cells were washed once with PBS, scraped, and lysed for 20 min on ice with M-PER^®^ Mammalian Protein Extraction Reagent (Thermo Scientific) supplemented with phosphatase inhibitors (Roche, Germany) and protease inhibitors (Roche, Germany). The lysate was then centrifuged for 20 min at 14 000 g, 4 °C. The supernatant was collected, and the protein concentration was quantified by a Bradford assay. Samples were prepared for SDS-PAGE in Laemmli buffer containing β-mercaptoethanol as a reducing agent. Subsequently, samples were processed in an SDS-PAGE (12% acrylamide) and transferred onto a nitrocellulose membrane (Bio-Rad, Germany). The membrane was then blocked with milk and probed with the specified antibodies. Developing was carried out using the Licor Odyssey^®^ system.

### Oligomerization and membrane translocation of MLKL

Cells were mechanically lysed by passing them 30 times through a 25G (0.5 × 25 mm) syringe needle in isolation buffer (250 mM sucrose, 5 mM Tris, 2 mM EDTA, pH 7.4). Subsequently, the samples were centrifuged at 14 000 g for 10 min at 4 °C. The resulting supernatant was then centrifuged at 100 000 g for 1 h at 4 °C. The supernatant obtained after this step was collected as the cytosolic fraction, while the pellet was washed with 500 µL of PBS and centrifuged again at 100 000 g for 1 h at 4 °C. The final pellet was collected as the membrane fraction. Both fractions were prepared for SDS-PAGE using the Criterion™ Cell system (4–15% gradient gel) (BIO-RAD) and analyzed by Western blotting as described above.

### TIRF imaging

After DOX and TSZ treatment were imaged in a widefield Elyra 7 SIM2 inverse, Carl Zeiss Microscope. Images were acquired in High Power-TIRF mode (HP-TIRF), with an alpha Plan-APOCHROMAT 63x/1.46 Oil objective, and detected with a sCMOS camera, pco.edge. Excitation of GFP fluorophores was performed with a 488 nm laser at 12% intensity and 30 ms exposure. Pictures were taken every 25 ms for a total of 150 – 300 frames. Cells were kept at 37 °C during imaging. The software used for imaging was ZEN (black edition).

### Single-particle tracking analysis

The recorded videos were converted to TIF format and processed for single-particle tracking analysis using the TrackIt (release 1.6 of March 2024) software developed by Kuhn et. al., 2021, operated in MATLAB R2024a. The spot detection was done with a threshold factor of 1.5, and the tracking used the u-track linear + random motion algorithm. The minimum track length was set to 10 frames, with 1 gap frame, and 2 frames as the minimum track length before a gap frame. For the MSD fitting analysis, the shortest track was set to 30 frames, with a power law function. Long tracks were defined by 3 survived dark periods.

### Stoichiometry analysis

#### Single-particle fluorescence brightness analysis

Single-particle fluorescence brightness analysis was performed to estimate the oligomeric state of MLKL-EGFP based on the fluorescence intensity of individual particles. The number of molecules per particle was inferred by comparing the fluorescence intensity, measured in photon counts, with that of a monomeric EGFP reference of known stoichiometry. Monomeric EGFP intensity was determined from lysates of cells expressing soluble EGFP that were imaged under the same acquisition conditions as the mouse and human cell samples (Supplementary figure 4A,B). The measured monomer intensity was used to calculate the expected fluorescence intensity of MLKL-EGFP oligomers (Supplementary figure 4C,D).

The experimental fluorescence intensity distributions of detected particles were fitted using a multi-Gaussian model, in which each Gaussian component represented a distinct oligomeric species. To account for the incomplete maturation of EGFP, fluorescence intensities were corrected assuming that 80% of EGFP molecules were fluorescent due to chromophore maturation efficiency (Schmidt, 1996; Subburaj *et al*., 2015). The relative abundance of each oligomeric species was subsequently calculated by correcting the measured occurrence according to the labeling efficiency and weighting each corrected occurrence by its corresponding oligomeric state (N-mer).

Image analysis was performed using the Stoichiometry Analysis Software (SAS) developed by Danial *et al*., 2022 and modified by Jonas Aufdermauer. The detection parameters were optimized as follows: ROI radius = 6 pixels, minimum sigma = 3, maximum sigma = 5, and detection threshold = 0.0002. Each set of videos corresponding to the different cell lines and experimental conditions was analyzed together with GFP lysate calibration samples acquired under identical imaging conditions. The labeling efficiency parameter was set to 80% for all analyses.

#### Monomeric EGFP calibration

The fluorescence intensity of monomeric EGFP was determined by single-molecule photobleaching analysis. Individual fluorescent particles were continuously illuminated while their fluorescence intensity was recorded over time. Because photobleaching occurs stochastically, individual bleaching events produced discrete decreases in fluorescence intensity that were assumed to correspond to the loss of a single EGFP fluorophore. Only particles classified as monomers were used to determine the mean fluorescence intensity and distribution of monomeric EGFP, which served as the calibration reference for the stoichiometry analysis.

#### Preparation of EGFP calibration samples

Calibration samples were generated by transient expression of monomeric EGFP (pEGFP-N2) in NIH-3T3 cells. Cells were trypsinized, counted, and seeded at a subconfluent density (<50% confluence). Twenty-four hours after seeding, cells were transfected with 200 ng pEGFP-N2 plasmid using Lipofectamine 2000 (Invitrogen, Germany) in Opti-MEM (Thermo Fisher Scientific, Germany), following the manufacturer’s recommendations. Cells were incubated for an additional 24 h before harvesting.

Cell pellets were lysed for 20 min at 4°C in lysis buffer containing 10 mM Tris-HCl (pH 7.5), 150 mM NaCl, 0.5 mM EDTA, and 0.5% Nonidet NP-40, supplemented with phosphatase inhibitors (Roche, Germany). Lysates were centrifuged at 15,000 × g for 20 min at 4°C, and the supernatant was collected. The clarified lysates were transferred to 8-well glass-bottom chambers (IBIDI, cat. no. 80827) and imaged using the same microscope settings and acquisition parameters as those used for the experimental samples.

#### Photobleaching analysis

Photobleaching analysis was performed as previously described (Subburaj *et al*., 2015). Briefly, the fluorescence intensity of individual particles was recorded during continuous illumination until complete photobleaching. The number of fluorophores per particle was estimated by counting the number of photobleaching steps in the fluorescence intensity trace.

Particle detection and fluorescence tracking were performed using the Stoichiometry Analysis Software (SAS) with the same detection parameters used for the fluorescence brightness analysis (ROI radius = 6 pixels, minimum sigma = 3, maximum sigma = 5, and detection threshold = 0.0002). The software automatically detected individual particles, tracked their fluorescence intensity over time, and quantified the number of discrete photobleaching steps for each particle. Particles were subsequently assigned to their corresponding oligomeric species according to the number of bleaching steps, and the percentage occurrence of each oligomeric species was calculated from the proportion of particles assigned to each class.

### Recombinant hMLKL1 expression and purification

Recombinant monomeric hMLKL1 was cloned into the pFastBac-GST-TEV vector, and purified as described in (Ros *et al*., 2025) Briefly, the resulting recombinant baculovirus was used to produce the protein in Sf9 insect cells. Cells were harvested 72 h post-infection and lysed by sonication in lysis buffer containing 25 mM HEPES (pH 7.5), 300 mM NaCl, 5% glycerol, 0.1% NP-40, and 5 mM β-mercaptoethanol, supplemented with complete protease inhibitor cocktail and DNase I. The supernatant from lysed cells was incubated with pre-equilibrated glutathione agarose beads (General Electric, Healthcare, Germany) at 4°C for 1 h with gentle agitation. Then, glutathione agarose beads with bound hMLKL1 were incubated with 50 μM TEV at 4 °C for 36 h. hMLKL1 was further purified using size-exclusion chromatography. The monomeric fraction was collected and verified by SDS-PAGE. Purified hMLKL1 was aliquoted and stored at −80 °C.

### Liposome permeabilization assays

Large unilamellar vesicles (LUVs) were prepared by mixing lipids (Avanti Polar Lipids: Alabaster, AL, USA) in chloroform at the molar ratios specified in Supplementary figure 6A to generate IPM1 (30% POPC, 40% DOPE, 25% DOPS, 5% PI), IPM2 (30% POPC, 40% DOPE, 25% DOPS, 5% PI(4,5)P₂), and IPM3 (30% POPC, 40% DOPE, 25% DOPS, 5% PI(3,4,5)P₃) compositions. The mixed lipids were evaporated under a nitrogen stream and dried under vacuum for 3 h. Lipid films were rehydrated in carboxyfluorescein buffer (80 mM carboxyfluorescein, 25 mM HEPES, and150 mM NaCl with pH 7.5) to a total lipid concentration of 2.5 mg/mL, subjected to 5 freeze–thaw cycles, and extruded 31 times through a 100 nm polycarbonate membrane (LiposoFast, Avestin). Unencapsulated dye was removed by size exclusion chromatography on a Sephadex G50 gravity column. LUVs were incubated with the indicated concentrations of hMLKL1 in assay buffer (25 mM HEPES and150 mM NaCl with pH 7.5) for 60 min at room temperature. Fluorescence was measured at (excitation 492 nm/emission 514 nm) using a TECAN plate reader. Maximum release was determined by adding 0.5% Triton X-100. Percentage release was calculated as (F_sample − F_baseline) / (F_max − F_baseline) × 100.

Giant unilamellar vesicles (GUVs) were generated from lipid mixtures (Avanti Polar Lipids: Alabaster, AL, USA) of POPC: Cardiolipin (8:2) and POPC: DOPS (8:2) containing 0.3% DiI (DiIC18(3); 1,1’-dioctadecyl-3,3,3’,3’-tetramethylindocarbocyanine). Lipids were solubilized in chloroform at 1.25 mg/mL and deposited onto platinum wire; the solvent was evaporated under vacuum for 30 min. GUVs were generated by electroformation in 300 mM sucrose buffer for 2 h. GUVs were then transferred to glass chambers pre-blotted with 5 mg/mL BSA for the permeabilization assay, performed in assay buffer (25 mM HEPES, 150 mM NaCl, 200 nM XDF-488, pH 7.5). hMLKL1 (200 nM) was added to GUVs, and images were acquired on a Zeiss LSM 980 confocal microscope. For two-color assays, a second dye (Alexa Fluor 655, 200 nM) was added to the imaging chamber following 75 min of incubation with hMLKL1 (200 nM). Images were analyzed using ImageJ. The kinetics of membrane permeabilization were quantified as the mean intravesicular fluorescence intensity normalized to the maximum signal.

### SLB and proteoliposome preparation for atomic force microscopy (AFM) imaging

Supported lipid bilayers (SLBs) were prepared by vesicle fusion on freshly cleaved mica. LUVs (20 µL, 2.5 mg/mL; fusion buffer: 25 mM HEPES pH 7.5, 150 mM NaCl, 2 mM CaCl₂) were deposited on mica and incubated for 30 min at 45 °C, then rinsed five times with imaging buffer (25 mM HEPES pH 7.5, 150 mM NaCl) to remove unfused vesicles and imaged immediately. hMLKL1 (200 nM) was added to pre-formed SLBs and incubated for 30 min at room temperature prior to imaging; control SLBs were treated with imaging buffer alone. Separately, hMLKL1 (200 nM) was incubated with LUVs (1 mg/mL) for 30 min at room temperature to form proteoliposomes. Proteoliposomes were deposited on mica, incubated for 15 min, and then washed prior to imaging.

### AFM imaging parameters

AFM imaging was performed in peak-force tapping mode (NanoWizard® ULTRA Speed 3, Bruker) in liquid at room temperature. SLBs and proteoliposomes were imaged using a cantilever (PEAKFORCE-HIRS-F-A; spring constant 0.35 N/m) at a scan rate of 2 kHz, with 512 × 512 pixel resolution over a 2 × 2 µm² scan area. Height images were flattened by first-order line-by-line subtraction. Protein clusters and membrane pores were identified and measured using JPKSPM Data Processing software (v8.1.31, Bruker). Pore diameters were estimated from the longest axis/equivalent circle diameter of the opening. Pore perimeter coverage (hole/MLKL ratio) was calculated as the fraction of the pore perimeter decorated by protein signal above background.

### Quantification and statistical analysis

Blots and images are representative of at least three independent experiments. All measurements were performed at least three times, and results are presented as mean ± SD. All statistical tests were carried out using GraphPad Prism 10 software. P-values from significantly different conditions are shown. Significance was calculated using an ordinary one-way ANOVA and a Tukey’s multiple comparisons test, otherwise it is specified in the figure legend.

### Image processing

The preprocessing of microscopy images was done using the Fiji (v 1.98) software. All images were assembled and edited using the Inkscape 1.3.2 software.

## Supporting information

Suppementary information

## Data availability

These data are available from the corresponding author upon reasonable request.

## Author contributions (CRediT)

V.M.-O. Conceptualization; Data curation; Formal analysis; Investigation; Methodology; Visualization; Writing-original draft. Q.Z. Formal analysis; Investigation; Methodology; Writing-review and editing. S.Z. Formal analysis; Investigation; Methodology; Writing-review and editing. J.A. Software; U.R. Conceptualization; Project administration; Supervision; Writing-original draft. A.G.-S. Conceptualization; Funding acquisition; Project administration; Supervision; Writing-original draft.

## Acknowledgements

We thank the CECAD Imaging Facility for its assistance. This work was funded by the German Research Foundation (CRC1403, grant ID 414786233; TRR353, grant ID 471011418; SPP2306 – project no. GA1641/7-1; CRC1507 –EXC-3094 – 533751785 to A.J.G.S.) and DFG-EXC 2030/2022 – 3017655 to the CECAD Imaging Facility. And by the Max Planck Society, as well as by the European Research Council (AdG MITOPORE 101195637).

We would like to thank all the technicians and facility staff for their support in this study: Annika Sarembe, Dr. Christian Jüngst, Julia Benecke, Lukas Faber, Christina Kunz, Anne Büttner, Tina Winkler, Sabine Wilhelm, and Gudrun Zimmer.

AI-assisted writing: OpenAI, GPT-5.5 and Claude (Sonnet 5) was used to improve the clarity, grammar, and organization of the manuscript. The AI was not used to generate data, perform analyses, or draw scientific conclusions. All content was reviewed and approved by the authors, who take full responsibility for the manuscript.

## References

Ando, Y. et al. (2020) ‘Necroptosis in pancreatic cancer promotes cancer cell migration and invasion by release of CXCL5’, PLOS ONE, 15(1), p. e0228015. Available at: 10.1371/journal.pone.0228015.

Bleicken, S. et al. (2018) ‘Topology of active, membrane-embedded Bax in the context of a toroidal pore’, Cell Death & Differentiation, 25(10), pp. 1717–1731. doi: 10.1038/s41418-018-0184-6.

Cai, Z. et al. (2014) ‘Plasma membrane translocation of trimerized MLKL protein is required for TNF-induced necroptosis’, Nature Cell Biology, 16(1), pp. 55–65. doi: 10.1038/ncb2883.

Chein, M., Perlson, E. and Roichman, Y. (2019) ‘Flow Arrest in the Plasma Membrane’, Biophysical Journal, 117(5), pp. 810–816. doi: 10.1016/j.bpj.2019.07.001.

Chen, X. et al. (2014) ‘Translocation of mixed lineage kinase domain-like protein to plasma membrane leads to necrotic cell death’, Cell Research, 24(1), pp. 105–121. doi: 10.1038/cr.2013.171.

Cho, Y. S. (2018) ‘The role of necroptosis in the treatment of diseases’, Frontiers in Pharmacology, 51(5), pp. 219–224. doi: 10.5483/BMBRep.2018.51.5.074.

Danial, J. S. H. et al. (2022) ‘Systematic Assessment of the Accuracy of Subunit Counting in Biomolecular Complexes Using Automated Single-Molecule Brightness Analysis’, Journal of Physical Chemistry Letters, 13(3), pp. 822–829. doi: 10.1021/acs.jpclett.1c03835.

Degen, M. et al. (2023) ‘Structural basis of NINJ1-mediated plasma membrane rupture in cell death’, Nature, 618(7967), pp. 1065–1071. doi: 10.1038/s41586-023-05991-z.

Fan, W. et al. (2019) ‘Flotillin-mediated endocytosis and ALIX–syntenin-1–mediated exocytosis protect the cell membrane from damage caused by necroptosis’, Science Signaling, 12(583). doi: 10.1126/scisignal.aaw3423.

Fish, K. N. (2022) ‘Total internal reflection fluorescence (TIRF) microscopy’, Current Protocols, 2(8), pp. 1–16. doi: 10.1002/cpz1.517.

Flores-Romero, H., Ros, U. and Garcia-Saez, A. J. (2020) ‘Pore formation in regulated cell death’, The EMBO Journal, 39(23), pp. 1–18. doi: 10.15252/embj.2020105753.

Garcia, L. R. et al. (2021) ‘Ubiquitylation of MLKL at lysine 219 positively regulates necroptosis-induced tissue injury and pathogen clearance’, Nature Communications, 12(1). doi: 10.1038/s41467-021-23474-5.

Garnish, S. E. et al. (2021) ‘Conformational interconversion of MLKL and disengagement from RIPK3 precede cell death by necroptosis’, Nature Communications, 12(1), p. 2211. doi: 10.1038/s41467-021-22400-z.

Gong, Y. N. et al. (2017) ‘ESCRT-III Acts Downstream of MLKL to Regulate Necroptotic Cell Death and Its Consequences’, Cell, 169(2), pp. 286–300.e16. doi: 10.1016/j.cell.2017.03.020.

Große, L. et al. (2016) ‘Bax assembles into large ring-like structures remodeling the mitochondrial outer membrane in apoptosis’, The EMBO Journal, 35(4), pp. 402–413.

Huang, D. et al. (2017) ‘The MLKL Channel in Necroptosis Is an Octamer Formed by Tetramers in a Dyadic Process’, Molecular and Cellular Biology, 37(5), pp. e00497–16. doi: 10.1128/mcb.00497-16.

Kuhn, T. et al. (2021) ‘Single molecule tracking and analysis framework including theory-predicted parameter settings’, Scientific Reports, 11(1), pp. 1–12. doi: 10.1038/s41598-021-88802-7.

Kulma, M. and Anderluh, G. (2021) ‘Beyond pore formation : reorganization of the plasma membrane induced by pore - forming proteins’, Cellular and Molecular Life Sciences, 78(17), pp. 6229–6249. doi: 10.1007/s00018-021-03914-7.

Lagerholm, B. C. et al. (2017) ‘Convergence of lateral dynamic measurements in the plasma membrane of live cells from single particle tracking and STED-FCS’, Journal of Physics D: Applied Physics, 50(6), p. 063001. doi: 10.1088/1361-6463/aa519e.

Li, L. et al. (2021) ‘The molecular mechanisms of MLKL-dependent and MLKL-independent necrosis’, Journal of molecular cell biology, 13(1), pp. 3–14.

Liu, S. et al. (2017) ‘MLKL forms disulfide bond-dependent amyloid-like polymers to induce necroptosis’, Proceedings of the National Academy of Sciences of the United States of America, 114(36), pp. E7450–E7459. doi: 10.1073/pnas.1707531114.

Liu, S. et al. (2024) ‘MLKL polymerization-induced lysosomal membrane permeabilization promotes necroptosis’, Cell Death and Differentiation, 31(1), pp. 40–52. doi: 10.1038/s41418-023-01237-7.

Martens, S. et al. (2021) ‘MLKL in cancer: more than a necroptosis regulator’, Cell Death and Differentiation, pp. 1757–1772. doi: 10.1038/s41418-021-00785-0.

Meng, Y. et al. (2023) ‘Phosphorylation-dependent pseudokinase domain dimerization drives full-length MLKL oligomerization’, Nature Communications, 14(1), pp. 1–18. doi: 10.1038/s41467-023-42255-w.

Moujalled, D., Strasser, A. and Liddell, J. R. (2021) ‘Molecular mechanisms of cell death in neurological diseases’, Cell Death and Differentiation, pp. 2029–2044. doi: 10.1038/s41418-021-00814-y.

Mulvihill, E. et al. (2018) ‘Mechanism of membrane pore formation by human gasdermin-D’, The EMBO Journal, 37(14), p. EMBJ201798321.

Noh, H. et al. (2026) ‘SIGLEC12 mediates plasma membrane rupture during necroptotic cell death’, Nature, 649(8096), pp. 460–466. doi: 10.1038/s41586-025-09741-1.

Oñate, M. et al. (2020) ‘The necroptosis machinery mediates axonal degeneration in a model of Parkinson disease’, Cell Death and Differentiation, 27(4), pp. 1169–1185. doi: 10.1038/s41418-019-0408-4.

Petrie, E. J. et al. (2018) ‘Conformational switching of the pseudokinase domain promotes human MLKL tetramerization and cell death by necroptosis’, Nature Communications, 9(1). doi: 10.1038/s41467-018-04714-7.

Petrie, E. J., Hildebrand, J. M. and Murphy, J. M. (2017) ‘Insane in the membrane: A structural perspective of MLKL function in necroptosis’, Immunology and Cell Biology, 95(2), pp. 152–159. doi: 10.1038/icb.2016.125.

Puylaert, P. et al. (2022) ‘Regulated Necrosis in Atherosclerosis’, *Arteriosclerosis*, Thrombosis, and Vascular Biology, 42(11), pp. 1283–1306. doi: 10.1161/ATVBAHA.122.318177.

Qian, S. et al. (2008) ‘Structure of transmembrane pore induced by Bax-derived peptide : Evidence for lipidic pores’, pnas, 105(45), pp. 17379–17383. doi: 10.1073/pnas.0807764105.

Richard, R. and Mousa, S. (2022) ‘Necroptosis in Alzheimer’s disease: Potential therapeutic target’, Biomedicine and Pharmacotherapy, 152, p. 113203. doi: 10.1016/j.biopha.2022.113203.

Ros, U. et al. (2017) ‘Necroptosis Execution is Mediated by Plasma Membrane Nanopores that are Independent of Calcium’, Cell Reports, 19(1), pp. 175–187. doi: 10.1016/j.celrep.2017.03.024.

Ros, U. et al. (2025) ‘MLKL activity requires a splicing-regulated, druggable intramolecular interaction’, Molecular Cell, 85(8), pp. 1589–1605. doi: 10.1016/j.molcel.2025.03.015.

Saha, S. et al. (2015) ‘Diffusion of GPI-anchored proteins is influenced by the activity of dynamic cortical actin’, Molecular Biology of the Cell, 26(22), pp. 4033–4045. doi: 10.1091/mbc.E15-06-0397.

Salvador-gallego, R. et al. (2016) ‘Bax assembly into rings and arcs in apoptotic mitochondria is linked to membrane pores’, 35(4), pp. 389–401.

Samson, A. L. et al. (2020) ‘MLKL trafficking and accumulation at the plasma membrane control the kinetics and threshold for necroptosis’, Nature Communications, 11(1), pp. 1–17. doi: 10.1038/s41467-020-16887-1.

Sandre, O., Moreaux, L. and Brochard-Wyart, F. (1999) ‘Dynamics of transient pores in stretched vesicles’, Proceedings of the National Academy of Sciences, 96(19), pp. 10591–10596. doi: 10.1073/pnas.96.19.10591.

Schmidt, T. (1996) ‘Local stoichiometries determined by counting individual molecules’, Analytical Chemistry, 68(24), pp. 4397–4401. doi: 10.1021/ac960710g.

Seifert, L. et al. (2016) ‘The necrosome promotes pancreatic oncogenesis via CXCL1 and Mincle-induced immune suppression’, Nature, 532(7598), pp. 245–249. doi: 10.1038/nature17403.

Subburaj, Y. et al. (2015) ‘Bax monomers form dimer units in the membrane that further self-assemble into multiple oligomeric species’, Nature Communications, 6(1), p. 8042. doi: 10.1038/ncomms9042.

Sudbrack, T. P. et al. (2011) ‘Observing the Solubilization of Lipid Bilayers by Detergents with Optical Microscopy of GUVs’, The Journal of Physical Chemistry B, 115(2), pp. 269–277.

Tamkun, M. M., O’Connell, K. M. S. and Rolig, A. S. (2007) ‘A cytoskeletal-based perimeter fence selectively corrals a sub-population of cell surface Kv2.1 channels’, Journal of Cell Science, 120(14), pp. 2413–2423. doi: 10.1242/jcs.007351.

Tanzer, M. C. et al. (2015) ‘Necroptosis signalling is tuned by phosphorylation of MLKL residues outside the pseudokinase domain activation loop’, Biochemical Journal, 471(2), pp. 255–265. doi: 10.1042/BJ20150678.

Tanzer, M. C. et al. (2016) ‘Evolutionary divergence of the necroptosis effector MLKL’, Cell Death and Differentiation, 23(7), pp. 1185–1197. doi: 10.1038/cdd.2015.169.

Wang, H. et al. (2014) ‘Mixed Lineage Kinase Domain-like Protein MLKL Causes Necrotic Membrane Disruption upon Phosphorylation by RIP3’, Molecular Cell, 54(1), pp. 133–146. doi: 10.1016/j.molcel.2014.03.003.

Wang, M. et al. (2025) ‘Amyloid Fibrillation of a Ninjurin-1-Derived α - Helical Peptide : Structural Insights into Conformational Transition’, ACS nano, 19(40), pp. 35977–35991. doi: 10.1021/acsnano.5c14731.

Xia, B. et al. (2016) ‘MLKL forms cation channels’, Cell Research, 26(5), pp. 517–528. doi: 10.1038/cr.2016.26.

Yang, L. et al. (2001) ‘Barrel-Stave Model or Toroidal ModelA Case Study on Melittin Pores’, Biophysical Journal, 81(3), pp. 1475–1485. doi: 10.1016/S0006-3495(01)75802-X.

Yoon, S. et al. (2017) ‘MLKL, the Protein that Mediates Necroptosis, Also Regulates Endosomal Trafficking and Extracellular Vesicle Generation’, Immunity, 47(1), pp. 51–65.e7. doi: 10.1016/j.immuni.2017.06.001.

Yu, Y. Q. et al. (2022) ‘The Role of Programmed Necrosis in Colorectal Cancer’, Cancers, 14(17), pp. 1–12. doi: 10.3390/cancers14174295.

Zacharias, D. A., Violin, J. D. and Newton, A. C. (2002) ‘Partitioning of Lipid-Modified Monomeric GFPs into Membrane Microdomains of Live Cells’, Science, 296(5569), pp. 913–916.

Zemel, A., Fattal, D. R. and Ben-shaul, A. (2003) ‘Energetics and Self-Assembly of Amphipathic Peptide Pores in Lipid Membranes’, Biophysical Journal, 84(4), pp. 2242–2255. doi: 10.1016/S0006-3495(03)75030-9.

