## Supplementary material for "Low-order MLKL oligomers induce irregular membrane defects in necroptosis": Suppementary information

### SUPPLEMENTARY FIGURES

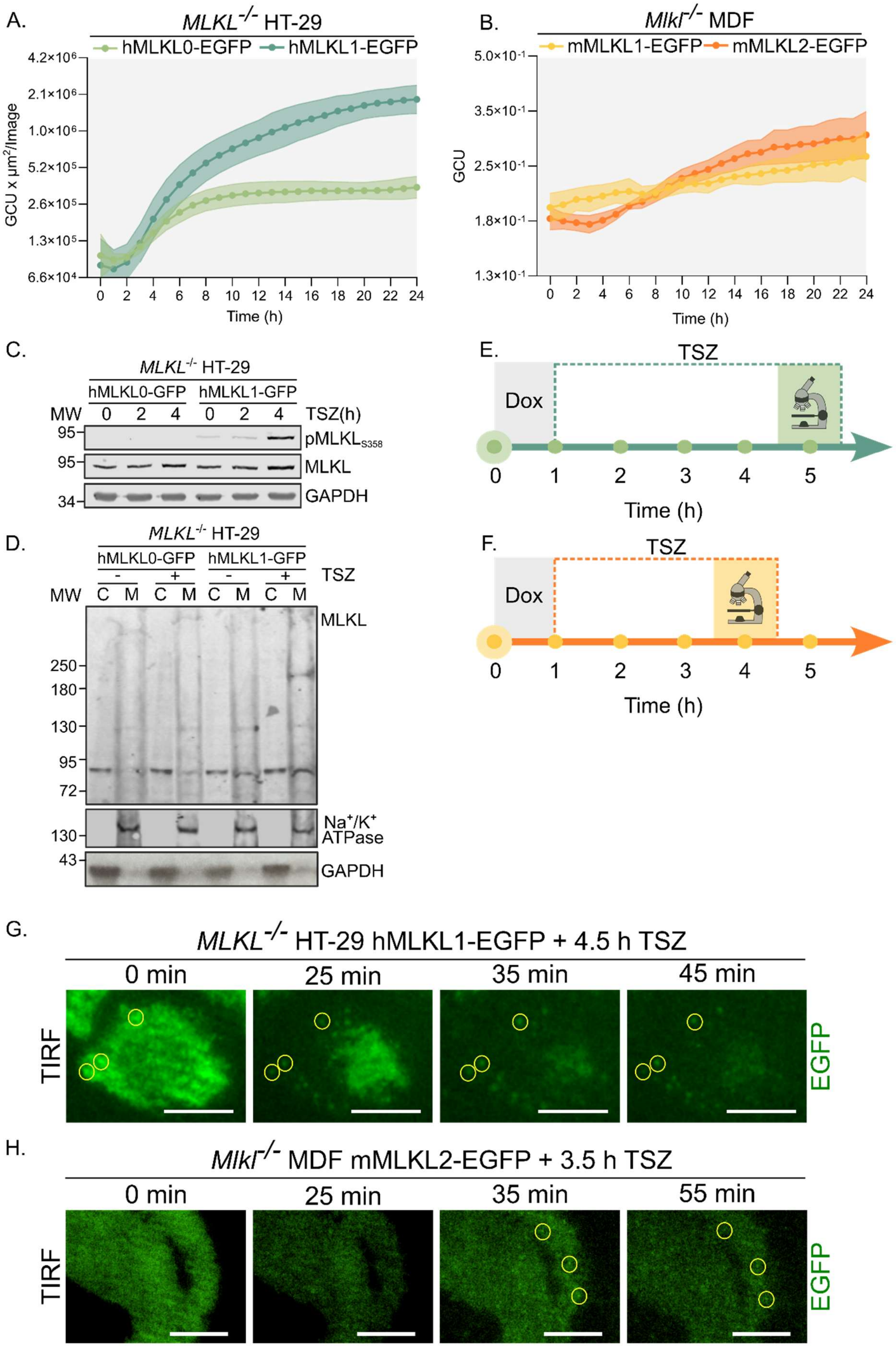

**Supplementary Figure 1. Characterization of MLKL-EGFP expression, phosphorylation, and membrane translocation in the Tet-On system.** (A, B) Total EGFP fluorescence intensity over time in *MLKL*$^{-/-}$ HT-29 (A) and *Mlkl*$^{-/-}$ MDF (B) cells expressing the indicated constructs following addition of doxycycline (Dox 1 μg/mL). The 4–5 h time window used for subsequent TIRF experiments is indicated by shading. Data are mean ± SD; n = 3 independent experiments. (C) Immunoblot for pMLKLS358 and total MLKL in *MLKL*$^{-/-}$ HT-29 cells expressing hMLKL0-EGFP or hMLKL1-EGFP, treated with TSZ for 0, 2, or 4 h. GAPDH serves as loading control. Phosphorylation at S358 is detected only in hMLKL1-expressing cells, confirming isoform-dependent RIPK3 substrate activity. (D) Subcellular fractionation and non-reducing SDS-PAGE of *MLKL*$^{-/-}$ HT-29 cells expressing hMLKL0-EGFP or hMLKL1-EGFP, untreated or TSZ-treated. C: cytosolic fraction; M: membrane fraction. MLKL is detected by anti-MLKL antibody; $Na^+/K^+$-ATPase and GAPDH serve as membrane and cytosolic fraction markers, respectively. hMLKL0 does not translocate to the membrane fraction and does not form high-molecular-weight complexes, providing a biochemical basis for its inactivity distinct from mMLKL1. (E) Experimental timeline for TIRF imaging of human cells: doxycycline addition at t = 0, TSZ treatment at t = 1 h, TIRF imaging window 4.5–5.5 h post-TSZ. (F) Experimental timeline for TIRF imaging of mouse MDF cells: Dox and TSZ co-treatment at t = 0, TIRF imaging window 3.5–4.5 h. (G) Higher-magnification view of the dashed white box region in Figure 1E, showing appearance of hMLKL1-EGFP particles (highlighted by yellow circles) at the plasma membrane during necroptosis. Scale bar, 10 μm. (H) As in (G) for Figure 1F (mMLKL2-EGFP in MDF cells). Scale bar, 10 μm.

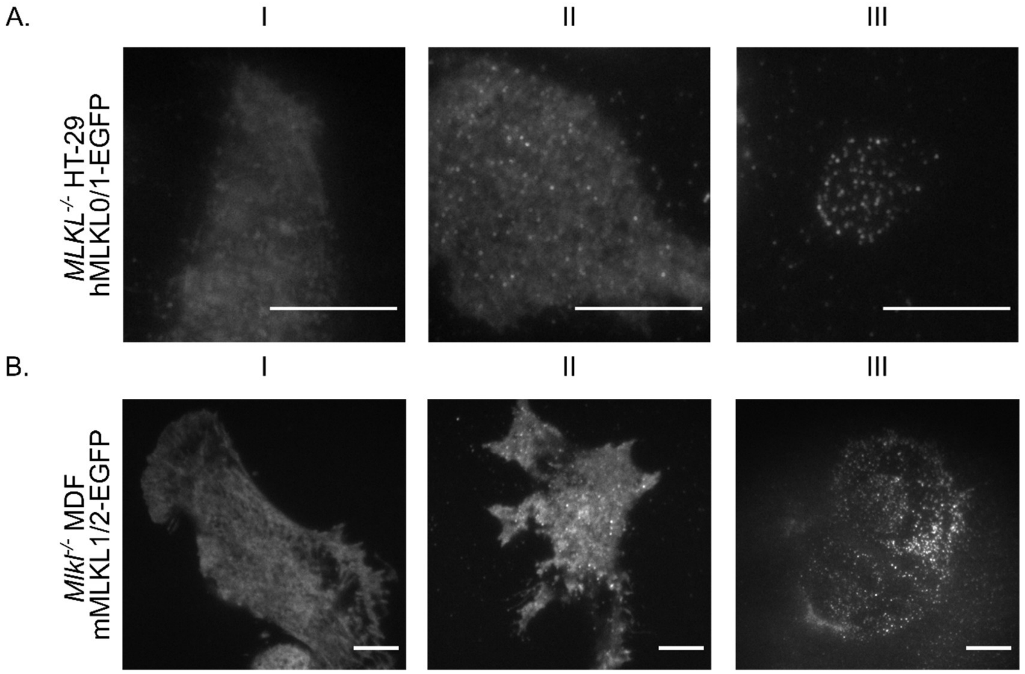


**Supplementary Figure 2. Representative TIRF images illustrating the three MLKL-EGFP distribution phenotypes.** (A) Representative TIRF images of *MLKL*$^{-/-}$ HT-29 cells expressing hMLKL0-EGFP or hMLKL1-EGFP, illustrating Type I (diffuse), Type II (diffuse + particles), and Type III (particles only) phenotype categories defined in Figure 2C. Scale bar, 10 µm. (B) As in (A) for *Mlkl*$^{-/-}$ MDF cells expressing mMLKL1-EGFP or mMLKL2-EGFP. Scale bar, 10 µm.

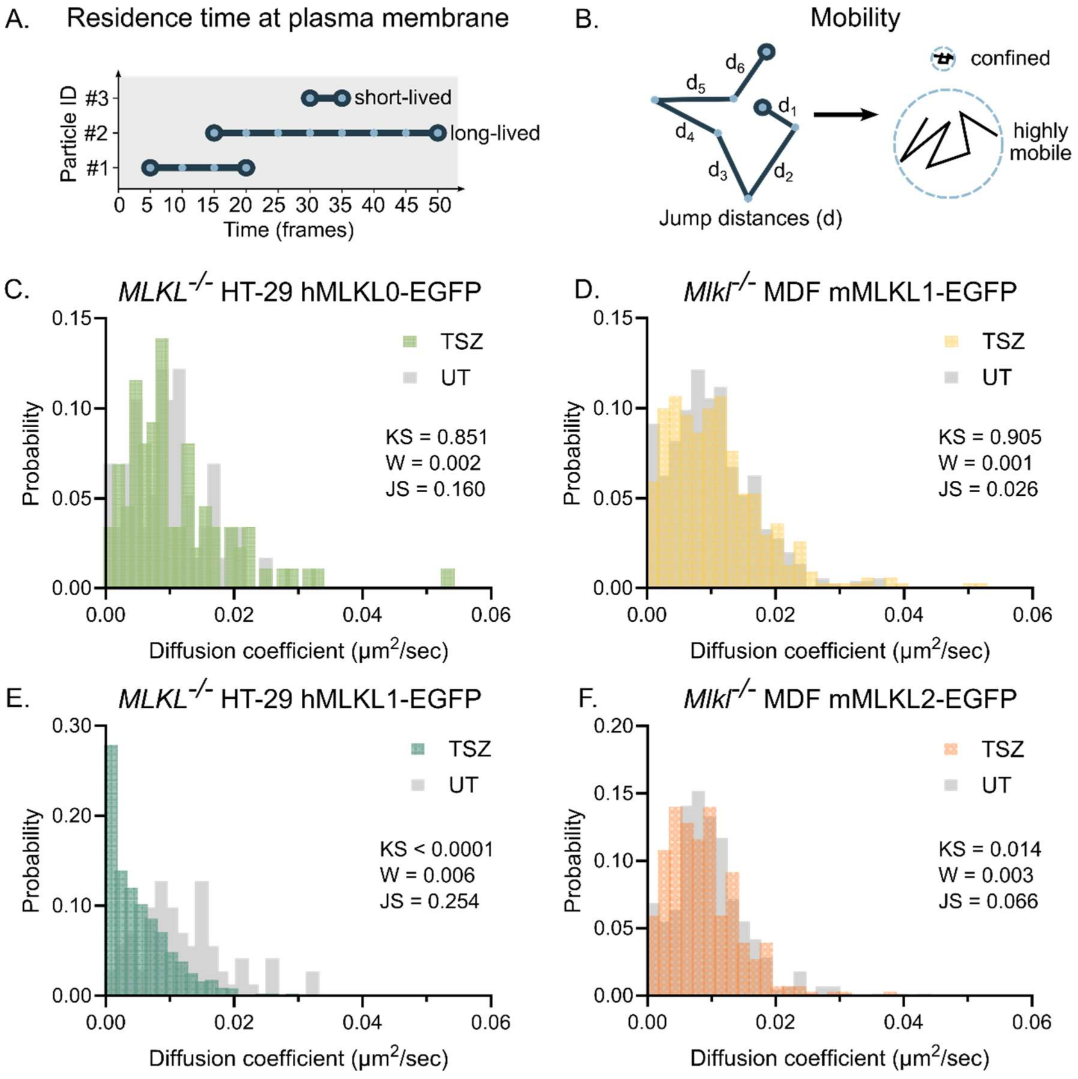


**Supplementary Figure 3. Tracking parameters and MSD-derived diffusion coefficients for all MLKL-EGFP isoforms.** (A) Residence time at the plasma membrane is reflected by the number of frames in which a particle is detected, thus, particles describing longer tracks are detected for a greater number of frames and are said to be long-lived particles, while particles with shorter tracks are considered short-lived. (B) The mobility of a particle is defined by the average of its jump distances (d) understood as the distance between the position of a particle in two consecutive frames. Confined particles are characterized for having smaller jump distances compared to mobile particles with greater jump distances. (C–F) Probability distributions of apparent diffusion coefficients estimated from mean squared displacement (MSD) analysis of individual particle tracks, for hMLKL0-EGFP (C), mMLKL1-EGFP (D), hMLKL1-EGFP (E), and mMLKL2-EGFP (F), under UT and TSZ conditions. Inset p-values indicate pairwise comparisons between UT and TSZ distributions: KS, Kolmogorov–Smirnov test statistic; W, Wasserstein distance; JS, Jensen–Shannon divergence. Only active isoforms (D, F) show statistically significant shifts in diffusion coefficient upon TSZ treatment, consistent with restricted membrane mobility. TSZ, necroptosis-inducing cocktail; UT, untreated; MSD, mean squared displacement.

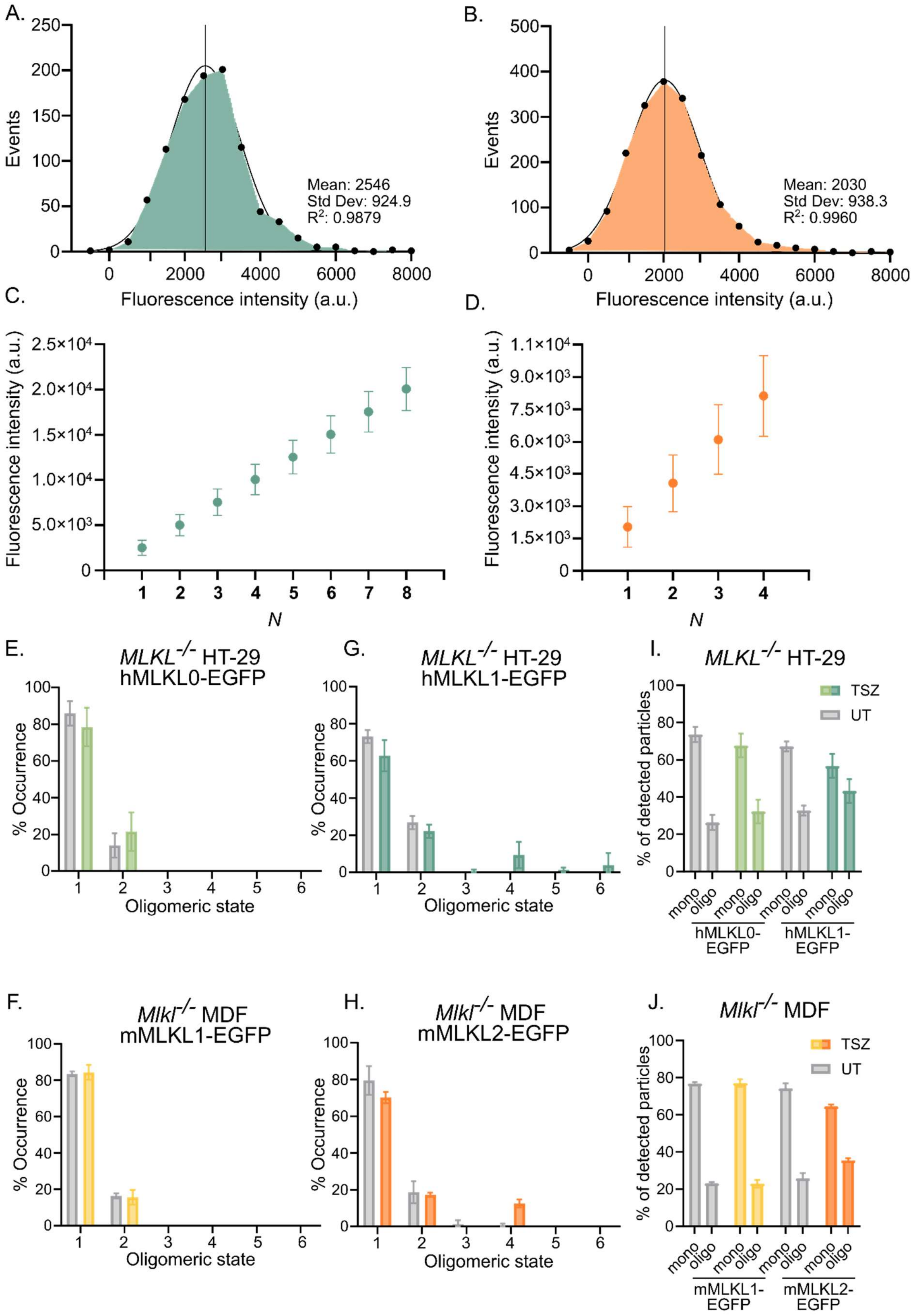
A.
Events
Mean: 2546
Std Dev: 924.9
R2: 0.9879
Fluorescence intensity (a.u.)
B.
Events
Mean: 2030
Std Dev: 938.3
R2: 0.9960
Fluorescence intensity (a.u.)
C.
Fluorescence intensity (a.u.)
N
D.
Fluorescence intensity (a.u.)
N
E.
MLKL-/- HT-29
hMLKL0-EGFP
% Occurrence
Oligomeric state
G.
MLKL-/- HT-29
hMLKL1-EGFP
% Occurrence
Oligomeric state
I.
MLKL-/- HT-29
TSZ
UT
% of detected particles
mono
oligo
hMLKL0-EGFP
hMLKL1-EGFP
F.
Mlkl-/- MDF
mMLKL1-EGFP
% Occurrence
Oligomeric state
H.
Mlkl-/- MDF
mMLKL2-EGFP
% Occurrence
Oligomeric state
J.
Mlkl-/- MDF
TSZ
UT
% of detected particles
mono
oligo
mMLKL1-EGFP
mMLKL2-EGFP

**Supplementary Figure 4. EGFP calibration and single-particle brightness stoichiometry data.** (A, B) Brightness distributions of soluble mEGFP from calibration lysates used for human cell experiments (A, mean intensity 2546 a.u., $R^2 = 0.988$) and mouse cell experiments (B, mean intensity 2030 a.u., $R^2 = 0.996$). The slightly different mean values reflect differences in TIRF imaging conditions between the two experimental sets; each set was independently calibrated. Gaussian fit shown as black line. (C, D) Expected fluorescence intensities (mean ± SD) for oligomeric species N = 1 through 8 (human, C) or N = 1 through 4 (mouse, D), calculated from the monomeric parameters in (A) and (B) respectively. Error bars propagate the SD of the monomer distribution. (E-H) Fractional occurrence of oligomeric states for all four isoforms in untreated and TSZ conditions: hMLKL0-EGFP (E), mMLKL1-EGFP (F), hMLKL1-EGFP (G), mMLKL2-EGFP (H). These panels show corrected occurrence values for 80% labelling efficiency. (I,J) Percentage of particles detected as monomers or oligomers in human (I) and mouse (J) cells. Data are mean ± SD; data from 3 independent experiments (n=3).

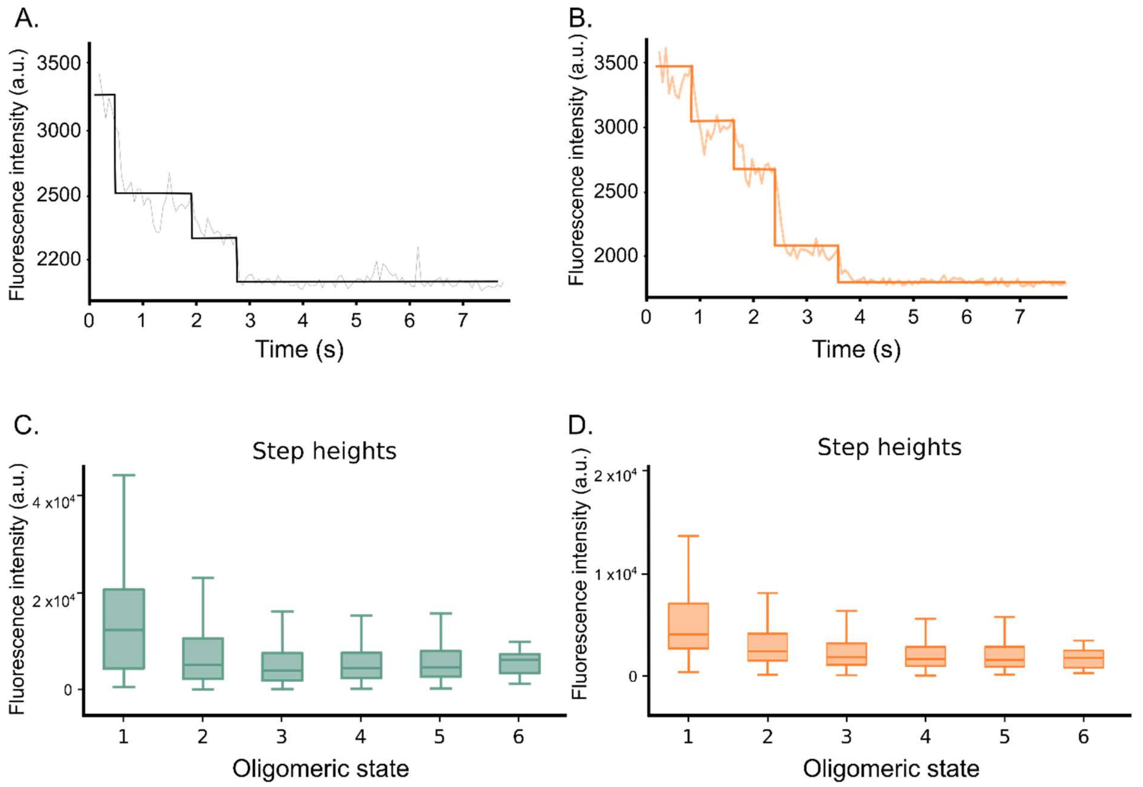


**Supplementary Figure 5. Photobleaching step profiles and linearity controls for stoichiometry analysis.** (A, B) Representative fluorescence intensity traces over time for individual MLKL-EGFP particles in hMLKL1-EGFP HT-29 cells (A) and mMLKL2-EGFP MDF cells (B) treated with TSZ. Stepwise decreases in intensity correspond to single EGFP photobleaching events. Fitted step profile shown as colored overlay. (C, D) Fluorescence intensity of individual photobleaching steps as a function of step number (i.e., bleaching event order within a trace), for hMLKL1-EGFP. Mean ± SD.

A.

| Name | Type | %POPC | %DOPE | %DOPS | %PI | %PI(4,5)$P_2$ | %PI(3,4,5)$P_3$ | %CL |
|---|---|---|---|---|---|---|---|---|
| IMP1 | Inner plasma membrane | 30 | 40 | 25 | 5 | 0 | 0 | 0 |
| IMP2 | Inner plasma membrane | 30 | 40 | 25 | 0 | 5 | 0 | 0 |
| IMP3 | Inner plasma membrane | 30 | 40 | 25 | 0 | 0 | 5 | 0 |

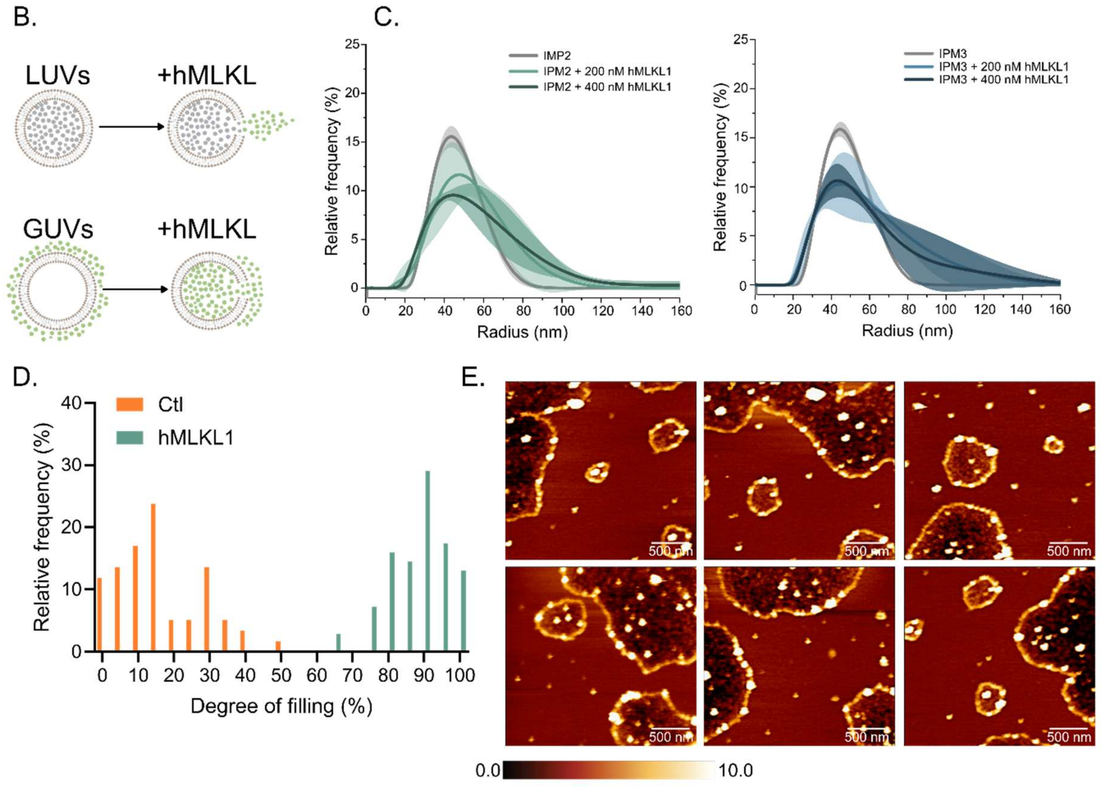


**Supplementary Figure 6. Lipid compositions, experimental strategy, and biophysical characterization of recombinant hMLKL1 in model membranes.** (A) Lipid compositions of the three inner plasma membrane mimetic LUV formulations used in this study (IPM1, IPM2, IPM3), differing in their phosphoinositide species (PI, PI(4,5)$P_2$, or PI(3,4,5)$P_3$ respectively; full molar percentages shown). (B) Schematic of LUVs and GUVs permeabilization assays. LUVs assay: carboxyfluorescein is encapsulated; protein addition causes dye release measured by fluorescence dequenching. GUVs assay: dye is present externally; protein addition causes dye internalization, measured by single-vesicle fluorescence microscopy. (C) Relative frequency distributions of LUVs hydrodynamic radius (by DLS) for IPM2 and IPM3 compositions, without and with addition of 200 or 400 nM hMLKL1. hMLKL1 addition broadens the radius distribution and shifts it toward larger values, consistent with protein-induced vesicle clustering or membrane remodeling. (D) Degree of individual GUVs filling in control (Ctl) samples (orange: no protein) compared to hMLKL1 treated samples (green). (E) Gallery of representative AFM images showing hMLKL1 clusters at SLBs edges, corresponding to the condition shown in Figure 6F (bottom). Six fields of view from 3 independent SLB preparations are shown. SLB, supported lipid bilayers; LUVs, large unilamellar vesicles; GUVs, giant unilamellar vesicles; AFM, atomic force microscopy; DLS, dynamic light scattering; IPM, inner plasma membrane.
